# Filling the Missing Puzzle: Integrating Interaction and Evolutionary Information to Enhance Protein Fitness Prediction in Inverse Folding

**DOI:** 10.64898/2026.10.05.756918

**Authors:** Zhihao Wang, Zhongyang Zhang, Yanmei Yang, Yuanyuan Qu, Yuguang Mu, Sheng Wang, Nanyu Han, Liangzhen Zheng, Weifeng Li

## Abstract

Protein inverse folding, which infers compatible amino acid sequences from a given backbone structure, is central to rational protein design. Current methods rely largely on sequence recovery rate as the evaluation metric, emphasizing structural compatibility but seldom addressing function. Yet functional activity depends on the proper network of non-covalent interactions—protein–ligand contacts being a prime example. We argue that function-oriented protein design is governed by three complementary constraints: structural, interaction, and evolutionary, none of which are handled in a unified manner by existing pretrained models. To bridge this gap, we introduce zPolaron, a graph neural network built on protein–ligand complexes for functional protein design. zPolaron represents residues and ligand atoms as a heterogeneous graph and jointly encodes structural, interaction, and evolutionary information by fusing physical-interaction-enhanced graph representations with ESM-2 embeddings. Across multiple fitness prediction benchmarks, zPolaron outperforms all baselines, with gains evident across structural, interaction, and evolutionary dimensions. Wet-lab experiments further confirm its utility in functional enzyme screening and design: the optimized CalA6-D205A-S160G mutant achieves a 17.84-fold improvement in activity, while the CalB-I121M mutant yields a 1.91-fold increase. Together, these results show that integrating interaction and evolutionary information fills the missing puzzle of inverse folding, enhancing protein fitness prediction beyond structural compatibility. zPolaron is available at https://github.com/zelixirSH/zPolaron.

## 1 Introduction

Proteins perform essential biological functions, including catalysis, molecular recognition, transport, and regulation, through their specific three-dimensional structures [1, 2, 3, 4, 5, 6, 7, 8]. Accordingly, protein design has long been regarded as an inverse inference problem from structure to sequence, aiming to identify amino acid sequences that can stably fold into a target structure while maintaining its function [9, 10, 11]. Inverse folding provides an important framework for structure-guided protein design. Given a protein backbone structure *X*, the objective of inverse folding is to search for an amino acid sequence *S* within the entire possible sequence space that is compatible with this structure[12, 13, 14, 15]. Early methods, represented by Rosetta [13], modeled this problem as energy optimization under fixed-backbone conditions. These approaches guided amino acid identity selection and side-chain conformational optimization through physical energy functions, achieving numerous milestones in the field of protein design [13, 16, 17, 18, 19]. Deep learning has established an alternative paradigm by directly learning the conditional probability distribution *P* (*S|X*) between structure and sequence from large-scale protein structure databases, leading to substantial improvements in inverse folding performance [14, 15, 20, 21, 22, 23, 24, 25, 26]. For instance, ProteinMPNN recovers native sequences with high accuracy using only local spatial neighborhood information [15]. ESM-IF further expanded the training scale by leveraging massive structural data predicted by AlphaFold [20]. Meanwhile, methods such as CarbonDesign explicitly model complex geometric relationships between residues through pairwise representations and triangular attention mechanisms [22]. Together, these studies demonstrate that models can effectively learn the constraints imposed by protein structures on sequences based on local packing and global topology.

Most existing inverse folding models employ the sequence recovery rate as their core optimization objective and evaluation metric, focusing essentially on structural compatibility. However, a high sequence recovery rate does not necessarily indicate successful protein design [27]. A mutation in even a single amino acid can cause a protein to misfold [5, 28]. Moreover, for a given protein structure, many different sequences can maintain the same folded state, whereas the native sequence represents only one outcome preserved under multiple evolutionary selection pressures rather than structural constraints alone [29, 2, 30]. In protein-ligand systems, functional activity is determined not only by the protein structure itself but also by whether the protein and ligand establish the correct network of non-covalent interactions. These interactions collectively determine substrate recognition, molecular binding, and catalytic activity [31, 32, 33]. For key residues within the binding pocket, amino acid selection must satisfy not only the requirements for local structural stability but also the chemical constraints required for ligand recognition and function [31, 34]. Such constraints may not be fully inferred from protein backbone geometry alone. Recent studies support this view. LigandMPNN was the first to explicitly incorporate ligand atoms as contextual information into the inverse folding framework, demonstrating the importance of non-protein components for function-oriented sequence design [21]. CARBonAra further enabled the modeling of different types of molecular entities using a geometric Transformer [35]. These studies suggest that incorporating the ligand environment, rather than relying solely on protein structural information, helps models learn constraints that are more directly related to protein function.

In addition to structural information, protein evolution also encodes rich functional constraints [36, 37, 38, 39]. Natural proteins have undergone billions of years of evolutionary selection, during which many functionally important residues have become highly conserved, while residues participating in the same structural or functional modules often exhibit complex co-evolutionary relationships. By learning from hundreds of millions of natural protein sequences, protein language models have been shown to implicitly capture sequence conservation, co-evolutionary patterns, and protein family-specific statistical regularities [40, 41, 42, 43, 44]. Furthermore, explicitly modeling protein-protein and protein-ligand interactions also provides valuable information. Previous studies have shown that knowledge-based models can achieve performance comparable to, or even surpassing, state-of-the-art deep learning methods in structure prediction and binding pose scoring [45, 46, 47]. Further analysis suggests that ligands with high affinity for the same binding pocket often exhibit similar interaction patterns [47]. Evolutionary constraints and protein-ligand interactions therefore not independent. Many highly conserved functional residues have been preserved throughout evolution precisely because they are responsible for essential ligand recognition or catalytic functions. Therefore, integrating backbone structure, evolutionary priors, and local physicochemical properties may be critical for function-oriented protein design.

Based on these observations, we propose that function-oriented protein design is fundamentally governed by three complementary types of constraints. Structural constraints determine whether a protein can fold correctly, interaction constraints determine whether it can accomplish molecular recognition and catalysis, and evolutionary constraints reflect functional preferences shaped by long-term natural selection. Existing inverse folding methods typically focus on one or two of these sources of information, whereas approaches that integrate structural information, protein-ligand interaction information, and evolutionary information within a unified framework remain limited. To address this gap, we present zPolaron, a graph neural network-based inverse folding model for protein-ligand complexes. In zPolaron, protein residues and ligand atoms are represented as different node types, while six types of heterogeneous edges explicitly encode geometric relationships and physicochemical interactions between residues and between residues and ligands. Evolutionary representations derived from ESM2 are incorporated as residue node features, and a confidence-guided iterative design strategy is introduced to jointly model structural information, interaction information, and evolutionary information. Unlike existing methods that mainly optimize sequence recovery, zPolaron aims not only to generate sequences compatible with the target structure but also to capture the key constraints underlying protein adaptability and function. To systematically evaluate this hypothesis, we assessed the model from multiple perspectives, including sequence recovery, evolutionary conservation, and fitness prediction. And wet-lab validation further confirms that zPolaron-guided enzyme discovery and mutation design yield substantial gains in catalytic activity. By integrating interaction and evolutionary information with structural modeling, zPolaron fills the missing puzzle of inverse folding and enhances protein fitness prediction during sequence design.

## 2 Methods

### 2.1 Model Architecture of zPolaron

#### 2.1.1 Heterogeneous Graph Framework and Message Passing Mechanism

Protein–ligand complexes contain rich heterogeneous information. zPolaron models the complex interaction patterns within the binding pocket more precisely by constructing a heterogeneous graph [48].

Let the heterogeneous graph be denoted as *G* = (*V, E*), where *V* = *V_r_ ∪ V_l_* comprises protein residue nodes *V_r_* and ligand atom nodes *V_l_*, and *E* = *E_rr_ ∪E_ll_ ∪E_rl_* comprises inter-residue edges, intra-ligand edges, and residue–ligand edges. Each message-passing block updates node hidden states as:

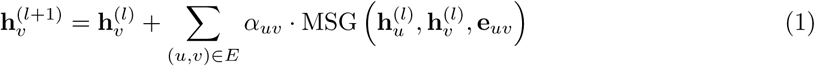

where **h**^(*l*)^ is the hidden state of node *v* after the *l*-th block, *α_uv_* is the attention weight, **e***_uv_* is the edge feature, and MSG is a learnable message generation function.

The construction of the heterogeneous graph fully accounts for the structural biology characteristics of the binding pocket. At the node level, the model distinguishes two heterogeneous node types: protein residue nodes and ligand atom nodes. Protein residue node features **x***_r_ ∈* R^667^ integrate evolutionary information from the pretrained language model ESM-2 (esm2 t30 150M UR50D, 150M parameters, 640-dimensional output from layer 30) and geometric information from the three-dimensional structure [40]. Ligand atom node features **x***_l_ ∈* R^24^ are represented by one-hot encoding of atom types. At the edge level, the model defines six distinct edge types *E* = *{E_rb_, E_rg_, E_ri_, E_lb_, E_lg_, E_li_}*, corresponding to different interaction patterns within the protein and between the protein and the ligand. Edges between protein residues include not only traditional peptide bond connections but also geometry-based edges defined by spatial proximity and interaction edges based on physicochemical properties. Edges between ligand atoms and protein residues encode various types of non-covalent interactions, such as hydrogen bonds, hydrophobic contacts, *π*–*π* stacking, etc. The interactions used here originate from the work of Ferreira de Freitas and Schapira [49]. Edge features are denoted as **e***_uv_ ∈* R*^de^* , where *d_e_* is the feature dimension of the corresponding edge type. This heterogeneous graph representation enables the model to adopt differentiated message-passing strategies for different node and edge types:

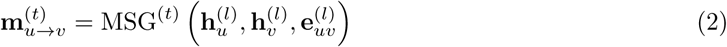

where *t* denotes different edge types and MSG^(*t*)^ is the specialized message function for the corresponding edge type.

#### 2.1.2 Confidence-Guided Iterative Design

zPolaron employs a confidence-guided iterative design strategy, a compromise between traditional auto-regressive generation and single-pass global prediction. Specifically, the model first adds mask tokens to all residues to be designed, then outputs the predicted probability distribution **p***_i_ ∈* R^20^ over the twenty natural amino acids at each masked position through the heterogeneous graph neural network. The maximum value in the predicted probability distribution reflects the model’s confidence in its prediction at that position:

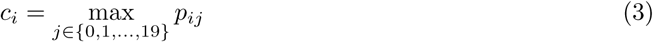

where *p_ij_* is the probability of the *j*-th amino acid at the *i*-th position in the model’s predicted probability distribution. After each iteration round, the model ranks the currently masked residues by confidence score and locks in the top fraction of residues with the highest confidence. The identities of the locked residues replace the original mask tokens and are re-input to the pretrained language model to obtain updated evolutionary context representations. This iterative strategy offers several advantages. First, it avoids the error accumulation problem of autoregressive methods, because only high-confidence residues are locked in each round. Second, it improves prediction accuracy by progressively introducing contextual information: as iteration proceeds, more residue identities become certain, providing richer context for the remaining masked residues. Third, it allows the model to adaptively allocate computational resources among residues of varying prediction difficulty—easily predicted residues are locked early, while difficult residues retain more rounds of optimization opportunity.

#### 2.1.3 Model Fitness Scoring

Beyond sequence design, zPolaron can also score a given protein sequence to evaluate its plausibility under a given structural context. Sequence scoring has important applications in protein engineering, such as zero-shot mutation effect prediction, protein fitness assessment, and deleterious mutation identification. zPolaron adopts a conditional-probability-based scoring strategy. For a given protein–ligand complex structure and a sequence to be evaluated *S* = (*s*_1_*, s*_2_*, . . . , s_L_*) *∈ {*0, 1*, . . . ,* 19*}^L^*, the model first inputs the sequence to the pretrained language model and graph neural network, then outputs the predicted probability distribution **p***_i_∈* R^20^ of the twenty natural amino acids at each position. The overall sequence score is defined as the mean negative log-likelihood across all positions [40, 50, 51]:

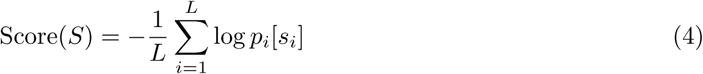

If the model consistently assigns high probabilities to the sequence, it indicates that the sequence is compatible with the given structural context and is therefore more likely to be physically plausible. A lower score indicates higher plausibility of the given sequence under the structural and evolutionary context. In addition to the overall negative log-likelihood score, the model can also compute per-residue confidence scores. For a given sequence *S*, the confidence at the *i*-th position is defined as the probability assigned to that amino acid:

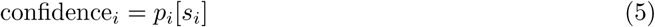

High confidence indicates that the model is highly certain about its prediction at that position, while low confidence suggests potential structural conflicts or mismatches in evolutionary pressure.

### 2.2 Datasets

#### 2.2.1 Dataset for Sequence Recovery

The training dataset originates from the protein–ligand complex dataset released by the LigandMPNN study [21]. This dataset contains protein–ligand complexes extracted from high-resolution crystal structures, totaling 62,908 samples. During data processing, random masking of protein sequences simulates the inverse folding task: given the structure and partial sequence information, the model predicts the original identities of masked residues. The validation set also comes from LigandMPNN’s protein–ligand complex validation set (3,515 samples), used to monitor generalization performance during training and to execute early stopping. The test set consists of 249 low-similarity PDB structures published between 2023 and 2025, collected independently. To ensure no significant sequence overlap between the test set and training set, we used MMseqs2 to align each protein chain of every test PDB against all chains of PDB entries released before 2023 [52]. A test PDB was included in the final test set only when its sequence similarity to any training sequence was strictly below 40%, thereby excluding structures with obvious homology to training data and ensuring the fairness and generalization capacity of the evaluation. The PDB entries comprising each of the dataset are listed in the Supplementary Materials.

#### 2.2.2 Dataset for Zero-Shot Scoring

For zero-shot scoring evaluation, we assembled datasets from multiple sources, covering three assessment tasks. The first task is protein stability. We drew PDB structures from the S2648 dataset and then applied three filters [53]. First, we kept only structures with more than 10 mutations to ensure statistically meaningful correlations. Second, we selected only single-chain structures, since ABACUS-T only supports single-chain input [26]. Third, we required the presence of at least one ligand, so that LigandMPNN and ProteinMPNN could be compared more fairly. After these steps, we obtained a final set of 10 PDB entries. The second task is protein fitness. Here we used eight enzyme targets from ProteinGym, all of which have clear annotations for both the enzyme identity and its substrate [50]. In addition, we included deep mutational scanning data for DAOx across five distinct substrates, taken from Vanella *et al* [54]. The third task is pocket disruption. For this, we used two datasets: the ADK dataset from Duncan *et al*., which contains enzymatic activity data for 178 adenylate kinase sequences from various species [55]; and the PETase dataset from Norton *et al*., which provides relative enzymatic activities for 214 PETase variants [56].

### 2.3 Model Training

The model is trained end-to-end in a multi-task learning framework. For each of the *N* training samples, the model outputs three quantities: node classification logits **z***_i_ ∈* R^20^, edge interaction counts ***µ****_i_ ∈* R*^de^* , and zero-inflation discrimination logits that distinguish structural zeros from count-generating zeros in the edge prediction task. The primary objective is node classification—predicting the amino acid identity of each protein residue from graph structure and contextual information— while the auxiliary edge prediction task estimates interaction-type counts between residue pairs and between residues and ligand atoms. The total loss combines these objectives as

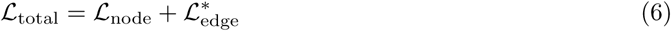

where *L*_node_ is the node classification loss and *L^∗^* denotes the edge prediction loss after applying dynamic weight scheduling and sparsity regularization (see Section 3 in Supplementary Materials for details).

Training used four NVIDIA A100 GPUs with distributed data parallelism. We employed the AdamW optimizer (*β*_1_ = 0.9, *β*_2_ = 0.98, *ɛ* = 1 *×* 10*^−^*^9^) with a weight decay of 0.01. The learning rate followed a warmup-plus-cosine-annealing schedule: linear warmup over the first 2,000 steps to the initial rate, followed by cosine decay to 1*×*10*^−^*^7^. Training was stopped when the validation node classification loss failed to improve by at least *δ* = 0.001 for 40 consecutive epochs. Key hyperparameters were: batch size 12 with gradient accumulation over 2 steps, initial learning rate 0.002, dropout rate 0.2, label distribution-aware smoothing with a base parameter of 0.05, and a maximum of 500 epochs. The edge prediction loss used a linear warmup over the first 10 epochs (maximum loss weight 8), with a sparsity regularization coefficient of 1 *×* 10*^−^*^5^.

### 2.4 Experimental Materials and Methods

#### 2.4.1 Genes, Strains, and Reagents

The genes encoding PsCalA, PsCalB, and their predicted novel enzyme variants were synthesized by Shanghai Zelixir Co., Ltd. All genes were cloned into the pET28a(+) vector [57], yielding recombinant plasmids pET28a-PsCalA and pET28a-PsCalB. Other novel enzymes discovered during the screening process were also ligated into the pET28a(+) vector.

*Escherichia coli* BL21(DE3) was used as the expression host (commercial strain). Isopropyl *β*-D-1-thiogalactopyranoside (IPTG), kanamycin (Kan), restriction endonuclease DpnI, and BeyoFusion™ Plus DNA polymerase were purchased from Beyotime (Shanghai, China). Protein quantification was performed using the BCA Protein Assay Kit (Thermo Fisher Scientific, USA). Substrates and cofactors including coniferyl alcohol, coniferaldehyde, and NAD^+^ were of analytical grade. The main instruments used in this study included a Shimadzu high-performance liquid chromatography (HPLC) system, a Huamei constant-temperature shaker, and a Xinzhi six-channel cell disruptor.

#### 2.4.2 Site-Directed Mutagenesis and Recombinant Plasmid Construction

Based on the residues identified through sequence and structural analyses, site-directed mutagenesis was performed via PCR using plasmid pET28a-PsCalA or pET28a-PsCalB as the template. The PCR program was as follows: 95 *^◦^*C for 30 s; 20 cycles of 95 *^◦^*C for 30 s, 55 *^◦^*C for 30 s, and 68 *^◦^*C for 3 min; followed by a final extension at 68 *^◦^*C for 10 min.

The PCR products were digested with DpnI to remove the template plasmid, and then transformed into *E. coli* BL21(DE3) competent cells. The transformation mixture was spread onto LB agar plates containing kanamycin (50 *µ*g*/*mL) and incubated overnight at 37 *^◦^*C to screen for positive clones.

#### 2.4.3 Protein Expression and Purification

A single colony was picked from the plate and inoculated into 4 mL of LB liquid medium containing kanamycin (50 *µ*g*/*mL), and cultured overnight at 37 *^◦^*C with shaking at 220 rpm. Then, 2 mL of the overnight culture was transferred into 100 mL of 2YT medium (supplemented with 50 *µ*g*/*mL kanamycin) at an inoculation volume of 2% (v/v). The culture was grown at 37 *^◦^*C and 220 rpm until the OD_600_ reached approximately 1.0–1.4. Protein expression was induced by adding IPTG to a final concentration of 0.2 mM, and the culture was further incubated at 20 *^◦^*C and 220 rpm for 16 h.

After induction, cells were harvested by centrifugation at 8,000 rpm for 5 min at 4 *^◦^*C. The cell pellet was resuspended in lysis buffer (pH 7.4) at a ratio of 1:5 (wet cell weight : buffer volume). The resuspension was sonicated using a cell disruptor for 15 min. The lysate was centrifuged at 12,000 rpm for 10 min at 4 *^◦^*C, and the supernatant was collected as crude enzyme extract.

Purification of the target enzymes was carried out by nickel-affinity chromatography on a Yonglian protein purification system. The buffers used were as follows: Lysis buffer: 20 mM Tris-HCl, 0.5 M NaCl, 10% (v/v) glycerol, pH 7.0; Elution buffer: 500 mM imidazole, 20 mM Tris-HCl, 0.5 M NaCl, 10% (v/v) glycerol, pH 7.0.

A 5 mL pre-packed Ni column was installed on the purification system with a flow rate set to 4 mL*/*min. The column was first equilibrated with lysis buffer until the UV baseline stabilized, after which the crude enzyme extract was loaded. After loading, the column was washed with a mixture of lysis buffer and elution buffer at a 15% ratio (elution buffer proportion) to remove non-specifically bound impurities. Once the UV signal returned to baseline, the target protein was eluted using 100% elution buffer, and the elution peak was collected. The eluate was subsequently concentrated using ultrafiltration centrifugal tubes, and the buffer was exchanged by repeated centrifugation with lysis buffer to thoroughly remove imidazole, yielding the purified target protein.

#### 2.4.4 Whole-Cell Catalysis Assays

Whole-cell catalysis assays were conducted using recombinant *E. coli* BL21(DE3) cells expressing the corresponding enzyme (without cell disruption or purification). After induction, cells were harvested and resuspended in PBS buffer (0.1 M, pH 7.4) at a ratio of 1 g wet cells per 5 mL buffer to prepare the cell suspension.

##### CalA whole-cell reaction system

Total volume 5 mL, containing 50 mmol*/*L Tris-HCl buffer (pH 9.0), 40 mmol*/*L coniferyl alcohol, 60 mmol*/*L NAD^+^, and 2.5% (v/v) of CalA cell suspension (i.e., 125 *µ*L). The reaction was carried out at 30 *^◦^*C for 1 h.

##### CalB whole-cell reaction system

Total volume 5 mL, containing 40 mmol*/*L Tris-HCl buffer (pH 9.0), 40 mmol*/*L coniferaldehyde, 60 mmol*/*L NAD^+^, and 2.5% (v/v) of CalB cell suspension (i.e., 125 *µ*L). The reaction was carried out at 30 *^◦^*C for 1 h.

After the reaction, 20 *µ*L of the reaction mixture was taken and mixed with 980 *µ*L of methanol to terminate the reaction and dilute the sample. The mixture was vortexed, centrifuged, filtered through a 0.22 *µ*m membrane, and analyzed by HPLC. Substrate consumption or product formation was quantified based on standard curves prepared using coniferyl alcohol/coniferaldehyde standards, thereby obtaining whole-cell catalytic activity data.

#### 2.4.5 Purified Enzyme Activity Assay

Purified enzyme activity was evaluated by monitoring the time course of the reaction using the target protein obtained after nickel-affinity chromatography. The purified enzyme solution was quantified by BCA assay and then diluted to a uniform protein concentration prior to the assay. Aliquots of 20 *µ*L were withdrawn from the reaction mixture every 5 min and immediately mixed with 980 *µ*L of methanol to terminate the reaction and dilute the sample. A total of six time points (0, 5, 10, 15, 20, and 25 min) were collected. Each sample was vortexed, centrifuged, filtered through a 0.22 *µ*m membrane, and analyzed by HPLC.

##### CalA reaction system

Total volume 5 mL, containing 50 mmol*/*L Tris-HCl buffer (pH 9.0), 40 mmol*/*L coniferyl alcohol, 60 mmol*/*L NAD^+^, and 5% (v/v) of diluted purified CalA enzyme solution. The reaction was carried out at 30 *^◦^*C.

Product concentrations at each time point were calculated based on a standard curve prepared using coniferaldehyde standards. The initial reaction rate was determined by linear regression of the linear portion of the product accumulation curve, where the slope represents the reaction velocity.

## 3 Results

zPolaron is a graph neural network model specifically designed for sequence design of protein-ligand complexes. As illustrated in Fig. 1, the model first represents protein residues and ligand atoms as a heterogeneous graph, so that the stacked message-passing blocks can pick up the intricate geometric and physicochemical interactions in the local microenvironment. The workflow proceeds in three steps: building the heterogeneous graph, encoding it through stacked message-passing blocks, and iteratively assigning the masked residues. After each iteration, a subset of high-confidence residues is fixed according to their confidence scores, and the predicted sequence is fed back into a pretrained protein language model to obtain richer evolutionary contextual information.

**Figure 1:**
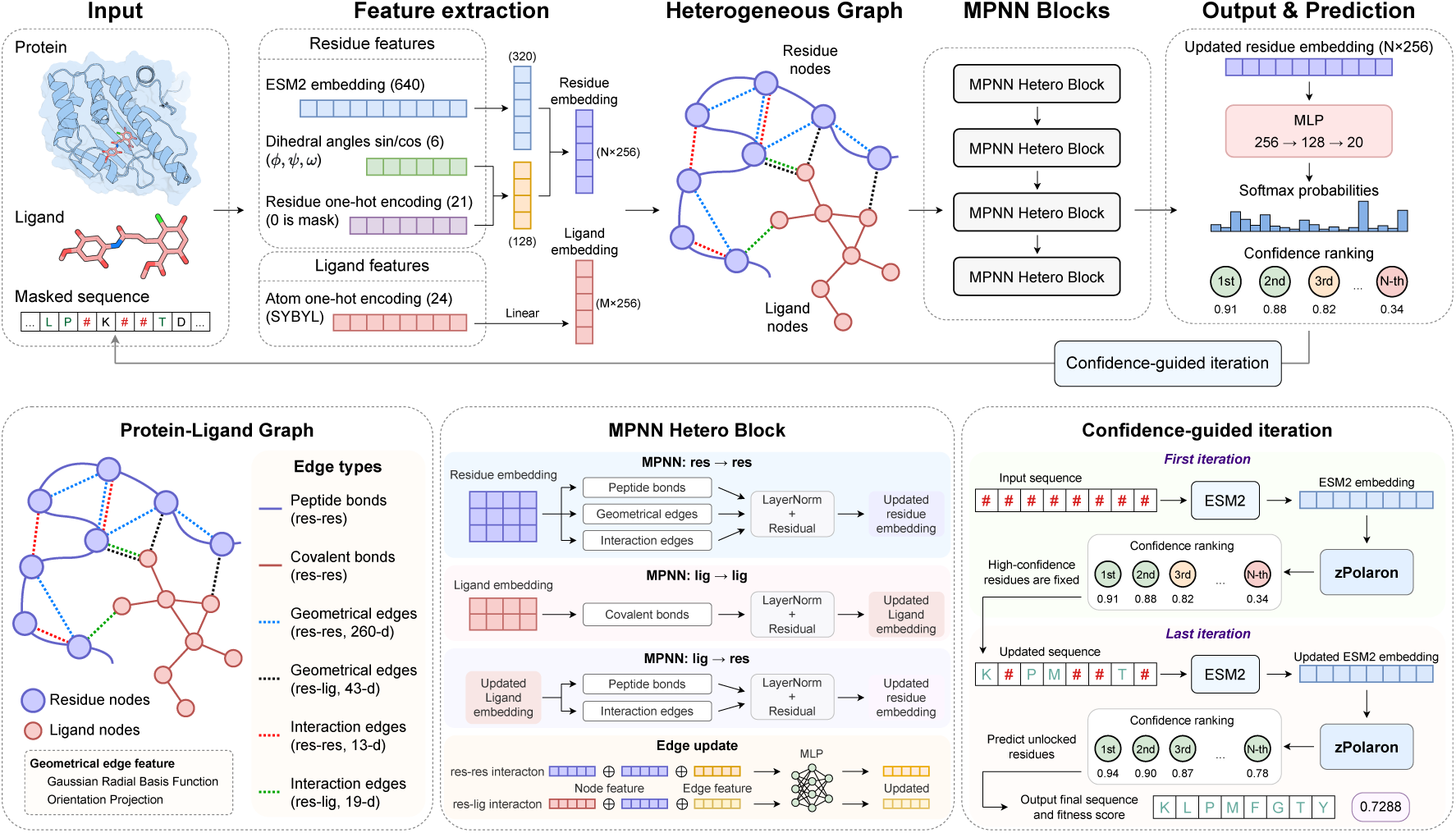
Schematic diagram of zPolaron.

### 3.1 zPolaron achieves the highest sequence recovery with strong consistency with evolutionary conservation

Structure-based sequence recovery is a standard evaluation method for inverse folding models, aiming to measure whether an algorithm can correctly infer the original amino acid sequence from a given protein structure. Therefore, we first evaluated the models on this metric. To avoid data leakage, this study collected PDB structures released between 2023 and 2025 as an independent test set. The sequence similarity between all test structures and the training data released before 2023 was strictly below 0.4. As shown in Fig. 2a, compared with current mainstream inverse folding models (including CARBonAra, ABACUS-T, LigandMPNN, and ProteinMPNN), zPolaron achieved a significantly higher sequence recovery rate. Furthermore, its overall recovery distribution is more concentrated, demonstrating its robustness when handling binding pockets with diverse properties.

**Figure 2:**
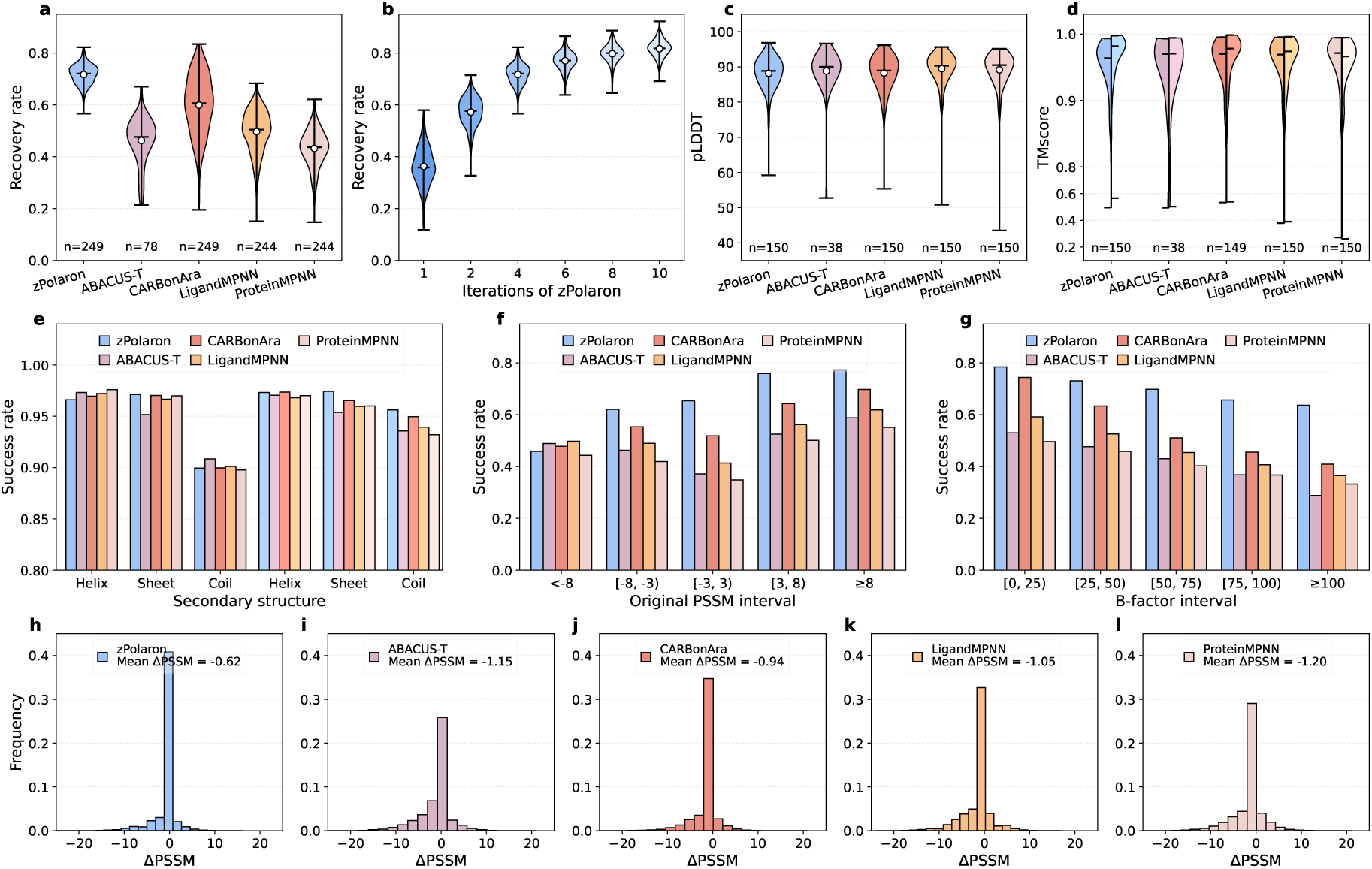
Sequence recovery and residue-level performance analysis of inverse folding models. (a) Comparison of sequence recovery among zPolaron and four baseline models. (b) Sequence recovery of zPolaron over successive confidence-guided design iterations. (c) Distribution of pLDDT scores for AlphaFold 3-predicted structures reconstructed from the designed sequences. (d) Structural similarity (TM-score) between the AlphaFold 3-predicted structures of the designed sequences and the input experimental structures (left half of each violin plot) and the AlphaFold 3-predicted structures of the corresponding native sequences (right half). (e) Residue-level secondary structure recovery. The left three groups of bars compare the designed structures with the experimental structures, whereas the right three groups compare them with the predicted structures of the native sequences. (f) Residue recovery rate as a function of the native residue PSSM score. (g) Residue recovery rate across different B-factor intervals. (h-l) Distributions of ΔPSSM, defined as the difference between the PSSM score of the predicted residue and that of the corresponding native residue.

Since zPolaron adopts a multi-round iterative design approach, Fig. 2b illustrates the detailed trend of its recovery rate across different iteration numbers. In the first iteration, because the initial sequence is completely masked, the pretrained language model ESM2 cannot capture effective sequence context. This leaves the initial embeddings lacking meaningful information (see the ablation study in the Supporting Materials for a detailed discussion), resulting in a low recovery rate and a wide distribution. However, with the introduction of the confidence-guided mechanism, high-confidence residues are progressively fixed to unlock the context, and the recovery rate exhibits a clear monotonic upward trend. By the 10th iteration, the median recovery rate exceeds 80%. Considering design accuracy, computational efficiency, and sequence diversity collectively, this study recommends using 4 iterations as the standard configuration.

To verify the structural feasibility of the designed sequences, we reconstructed their structures using AlphaFold 3 [46]. The results show that the sequences designed by all models exhibit extremely high structural confidence upon reconstruction, with pLDDT values close to 90 (Fig. 2c). We also calculated the TM-score [58, 59], a backbone similarity metric, between this reconstructed structure and both the input crystal structure and the AlphaFold 3-predicted structure of the native sequence (AF3 structure). Fig. 2d indicates that the TM-score for each method remains above 0.95, showing that all methods can well preserve the backbone topology of native proteins. It can be observed from the Fig. 2d that the predicted structures of the sequences designed by zPolaron resemble the predicted structures of the native sequences more than the crystal structures (the right half of each violin plot shows a higher average TM-score). A similar phenomenon occurs for CARBonAra, which has the second-highest sequence recovery rate. This phenomenon may stem from the prediction bias of AlphaFold 3 itself. Because output sequences with high recovery rates are closer to the native sequence, the structures of the new sequences predicted by AlphaFold 3 will also bias toward the AF3 structure. In terms of secondary structure (Fig. 2e), the designed sequences also closer resemble the predicted structure of the native sequence (the three clusters of bars on the right) rather than the crystal structure (the three clusters of bars on the left), especially for random coils. zPolaron demonstrates a strong capacity to recover *β*-sheet secondary structures. This may be because helical structures are primarily sustained by backbone hydrogen bonds, whereas *β*-strands can form a richer variety of interactions, such as side-chain *π*-*π* stacking or salt bridges, in addition to backbone hydrogen bonds. Consequently, zPolaron, which explicitly models interactions, can extract effective information more easily.

To thoroughly analyze the prediction preferences of the model for binding pocket residues, we evaluated the models across multiple dimensions at the amino acid level. We analyzed the relationship between residue recovery success rate and the Position-Specific Scoring Matrix (PSSM) of native residues. As shown in Fig. 2f, as the PSSM values of native residues increase, the recovery success rate of zPolaron exhibits a clear monotonic upward trend. In contrast, other baseline methods do not show this monotonic increasing feature strongly correlated with evolutionary conservation. We utilized ΔPSSM (the difference in PSSM between the output residue and the input residue) to quantify residue conservation, where ΔPSSM *>* 0 indicates that the mutation aligns better with the conservation preferences of native homologous sequences. The results in Figs. 2h-l show that although the sequences predicted by all inverse folding methods generally tend to shift ΔPSSM toward negative values, zPolaron achieves the highest average ΔPSSM (Mean ΔPSSM = *−*0.62), outperforming CARBonAra (*−*0.94), LigandMPNN (*−*1.05), ABACUS-T (*−*1.15), and ProteinMPNN (*−*1.20). This indicates that zPolaron possesses a superior capability to recover positions that are more conserved in native homologous sequences, demonstrating a stronger consistency between the model output and evolutionary conservation. It is worth noting that CARBonAra also predicts a residue-wise PSSM during inference and preferentially selects residues with higher predicted PSSM scores, which likely contributes to its second-best average ΔPSSM.

Fig. 2g illustrates the sensitivity of the model recovery success rate to the local disorder of the input structure, which is approximated by the B-factor. In the extremely stable, low B-factor interval [0, 25), all methods maintain high success rates. However, as structural uncertainty increases, the success rates of all methods drop noticeably. In comparison, zPolaron consistently maintains the highest success rate across different B-factor intervals, with a slower rate of decline. Ablation studies (Fig. S1) indicate that the model omitting backbone structural features yields the lowest recovery rate, failing to achieve a significant increase even with multiple iterations. This underscores that the backbone structure is the core component of inverse folding models. When the model incorporates backbone information while integrating either ESM embeddings or interaction information, the success rates across all intervals increase. Therefore, we infer that when the B-factor of the input structure is high (i.e., when three-dimensional coordinates exhibit uncertainty or high flexibility), purely geometry-based models like ProteinMPNN might be more susceptible to coordinate noise. Conversely, the sequence context provided by ESM2 evolutionary embeddings may partially compensate for the uncertainty in structural information. Although interaction features depend on geometry, they are typically encoded discretely based on distance thresholds or local geometric rules. Consequently, they may be more robust than pure coordinate representations to some extent, thereby helping the model mitigate the impact of structural noise. In practical applications, many drug targets of high therapeutic value, such as GPCRs [60, 61], ion channels [62, 63], and enzyme active centers with prominent conformational dynamics [64, 65], often feature functional pockets accompanied by high flexibility or local disorder, or sometimes must rely on poor-quality, low-resolution crystal structures. The insensitivity of zPolaron to structural uncertainty provides a distinct advantage when addressing these challenges.

To further evaluate the generalizability of zPolaron, we additionally tested it on a cytochrome P450 (CYP) dataset using structures predicted by AlphaFold 3 rather than experimentally determined structures [66, 67]. As shown in Fig. S3, the pretrained model maintained high sequence recovery on the predicted structures, and sequence recovery further improved after lightweight fine-tuning of only the output classification layer. These results suggest that zPolaron can be directly applied to AlphaFold3-generated structures and can be efficiently adapted to specific protein families with minimal retraining.

### 3.2 Conditional sequence distributions learned by inverse folding implicitly encode stability information

Early protein design methods, represented by Rosetta, optimized physics-based energy functions by selecting amino acid identities and optimizing side-chain conformations to reduce the free energy of a protein structure, thereby generating sequences compatible with a given backbone. From this perspective, these methods can be regarded as design strategies that optimize energy, or more specifically, protein stability. Deep learning-based protein design methods have also been reported to improve protein stability. Previous studies have shown that sequences designed by ProteinMPNN exhibit higher thermal stability than native protein sequence [15, 68]. Sun *et al.* also demonstrated that ESM-IF and ABACUS-R possess the capability for zero-shot prediction of ΔΔ*G* [69]. Therefore, in addition to the sequence recovery rate, we evaluated this metric as well. We employed the Mean Negative Log-Likelihood (Mean NLL) of the predicted probabilities across all residues as the scoring metric. For a fair comparison, the predicted probabilities of the other three baseline methods involved in the evaluation were also transformed using the aforementioned formula to output scores for the mutant sequences. CARBonAra adopts a different design strategy by generating sequences based on predicted PSSM values [35]. Therefore, its built-in sequence scoring function was used directly. Under this scoring scheme, all inverse folding methods based on mean negative log-likelihood are expected to show a negative correlation with the experimental measurements, whereas CARBonAra is expected to show a positive correlation. In other words, a lower mean negative log-likelihood indicates that a mutation is more favorable from both energetic and structural perspectives.

Figs. 3a and b present the Pearson and Spearman correlation matrices of all methods on the ten protein targets in the S2648 dataset [53]. ProteinMPNN achieved the best overall performance for thermostability prediction and obtained the highest correlation on several targets. However, its performance dropped sharply on specific proteins, such as 1AMQ and 5DFR. A similar trend was also observed for LigandMPNN. In contrast, although zPolaron did not consistently achieve the highest correlation across all targets, its Spearman correlation fell below 0.3 only on 2NVH, while remaining above 0.3 for all other proteins. These results indicate that zPolaron exhibits better generalization across different protein targets.

**Figure 3:**
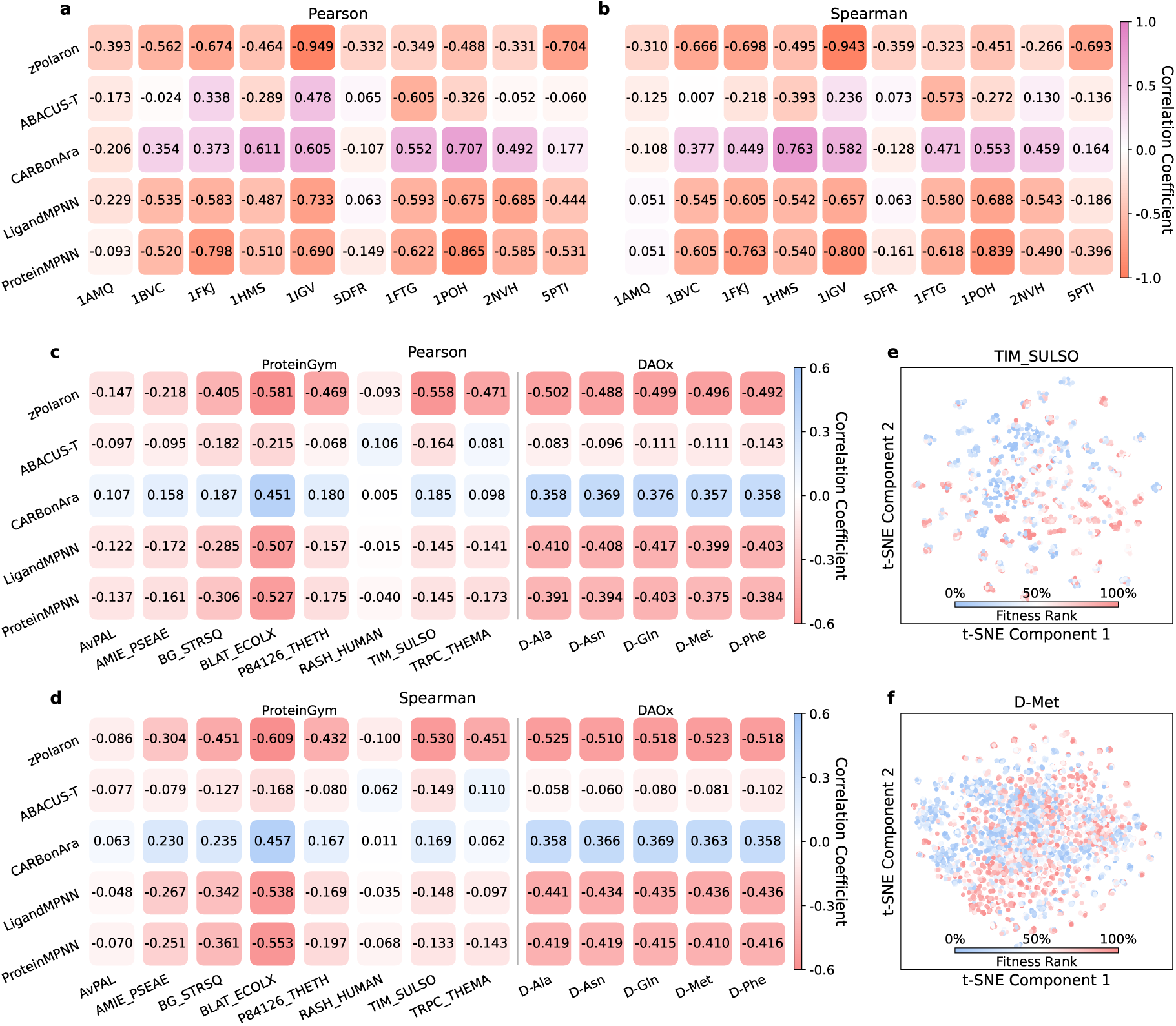
Correlation analysis of zero-shot scoring. (a, b) Pearson (a) and Spearman (b) correlations between zero-shot scores and ΔΔ*G* values on the S2648 protein stability dataset. (c, d) Pearson (c) and Spearman (d) correlations between zero-shot scores and experimentally measured fitness values on protein fitness benchmark datasets. (e) t-SNE visualization of the final embedding space of zPolaron for the TIM SULSO target. Data points are colored according to their experimentally measured fitness ranking. (f) t-SNE visualization of the final embedding space of zPolaron for the DAOx target with D-Met as the substrate.

### 3.3 Integrating three constraints enables superior zero-shot fitness prediction with strong practical utility

Sequence recovery and stability prediction primarily evaluate whether an inverse folding model can identify structurally compatible amino acids for a given protein backbone. However, the evolution of natural proteins is governed not only by structural constraints but also by functional requirements and evolutionary selection. A more stringent evaluation is therefore to examine whether the sequence probabilities generated by an inverse folding model can predict protein fitness [36, 38, 51]. Such an assessment provides insight into whether the learned representations extend beyond the sequence-structure relationship to capture the deeper coupling among sequence, structure, and function. To this end, we further investigated the relationship between model scores and experimentally measured fitness landscapes.

We evaluated both Pearson and Spearman correlations on two benchmark settings. The first consisted of eight enzyme systems with known substrates from the ProteinGym database, representing cross-target evaluation [50]. The second consisted of a DAOx enzyme system with five different substrates [54]. Performance across different targets reflects the model’s generalization ability and its adaptability to diverse structural topologies. Different enzyme targets possess distinct structural characteristics, including overall fold topology, catalytic residue arrangements, and electrostatic distributions. Good performance under this setting suggests that the model may capture underlying physicochemical and evolutionary principles, including backbone geometric constraints, hydrophobic core packing, and feasible residue dihedral conformations. Meanwhile, the correlation across different substrates for the same target reflects the substrate specificity of the model and its sensitivity to the non-covalent chemical microenvironment, verifying whether the model is capable to capture protein-ligand coupling.

Figs. 3c and d summarize the correlation results of all methods across the two evaluation settings. Overall, zPolaron achieved superior performance in most test systems. Across the eight ProteinGym enzyme targets, zPolaron consistently produced stronger negative correlations than the baseline methods, with the highest Spearman correlation coefficients reaching *−*0.609 and *−*0.530. In the DAOx multi-substrate benchmark, the correlation coefficients of zPolaron remained consistently close to *−*0.5, showing both relatively high absolute values and good consistency across different substrates. As representative and widely used inverse folding models, ProteinMPNN and LigandMPNN achieved moderate prediction performance. For example, their Spearman correlation coefficients on BLAT ECOLX reached *−*0.553 and *−*0.538, respectively. However, their performance varied substantially across different targets, exhibiting weak correlations for most enzyme systems. Furthermore, their correlations on the DAOx target across different substrates were also inferior to those of zPolaron. CARBonAra generally produced relatively weak positive correlations, with most values below 0.25 and only BLAT ECOLX exceeding 0.45, indicating that its generalized scoring capability is relatively limited. ABACUS-T showed relatively poor performance on this task. Even on BLAT ECOLX, where the other methods achieved relatively strong correlations, its correlation coefficient was only around *−*0.2. It also showed almost no correlation on the DAOx benchmark, suggesting that this method is better suited for sequence design than for sequence scoring. Ablation studies demonstrate that the ESM embeddings, backbone structural information, and interaction information in zPolaron all contribute to zero-shot prediction. Any combination of two components yields higher accuracy than the other four baseline models. Among them, the ESM information contributes the most to fitness scoring, highlighting the importance of protein language models with rich sequence knowledge for fitness-oriented evolutionary prediction.

To further investigate the learned representations, we extracted the embeddings from the final hidden layer and visualized them using t-SNE. As shown in Fig. 3e and f, the learned feature space spontaneously formed several dense local clusters. Moreover, these clusters exhibited a clear relationship with the experimentally measured fitness ranking, ranging from the highest-fitness variants (top 0%) to the lowest-fitness variants (bottom 100%). Variants with high and low fitness tended to occupy different regions of the latent space, although this pattern became less apparent for variants containing a large number of mutations because of the high density of data points. These results indicate that the learned latent representations contain information associated with experimental fitness and may capture an underlying mapping among sequence, structure, and function. This observation further suggests that the three-dimensional residue relationships and sequence contextual information learned by the graph neural network enable the predicted conditional probabilities to reflect the effects of mutations on structure- and function-related constraints.

### 3.4 zPolaron sensitively identifies loss-of-function mutations in binding pockets

Zero-shot fitness prediction reflects a model’s ability to capture the overall fitness landscape. While this type of correlation analysis validates the statistical association between model predictions and experimental phenotypes and do not directly reveal whether the model has learned the key structural constraints underlying protein function. In particular, mutations at critical residues within the binding pocket often simultaneously affect local structure, substrate recognition, and the catalytic microenvironment, providing a more direct test of the model’s ability to capture functional perception capability. Following the strategy of Masters *et al.* [70], we designed three extreme disruptive mutation modes for two enzyme systems, PETase (polyethylene terephthalate hydrolase) [56] and ADK (adenylate kinase) [55], to simulate the physicochemical collapse of the catalytic pocket. These modes include side-chain erasure (mutation to GLY), hydrophobic packing (mutation to PHE), and property inversion (mutation to a residue with completely opposite properties).

As shown in Figs. 4a-h, these mutations induce substantial structural changes in the key pocket regions of both enzymes. We mixed these negative variants, which are expected to completely lose catalytic function, with the corresponding wild-type libraries and evaluated each model’s ability to distinguish them using fitness scores. The ROC and precision-recall (PR) curves are presented in Figs. 4i-l. Among all methods, zPolaron showed the highest sensitivity to disruptions of the binding pocket microenvironment. On the PETase dataset, it achieved an AUC of 0.934 and an average precision (AP) of 0.847. On the ADK dataset, the performance further improved to an AUC of 0.995 and an AP of 0.988, indicating that the model was able to identify nearly all loss-of-function variants. In comparison, although all baseline methods performed better than random classification (AUC = 0.5 and AP = 0.25), their overall discriminative ability was substantially lower than that of zPolaron.

**Figure 4:**
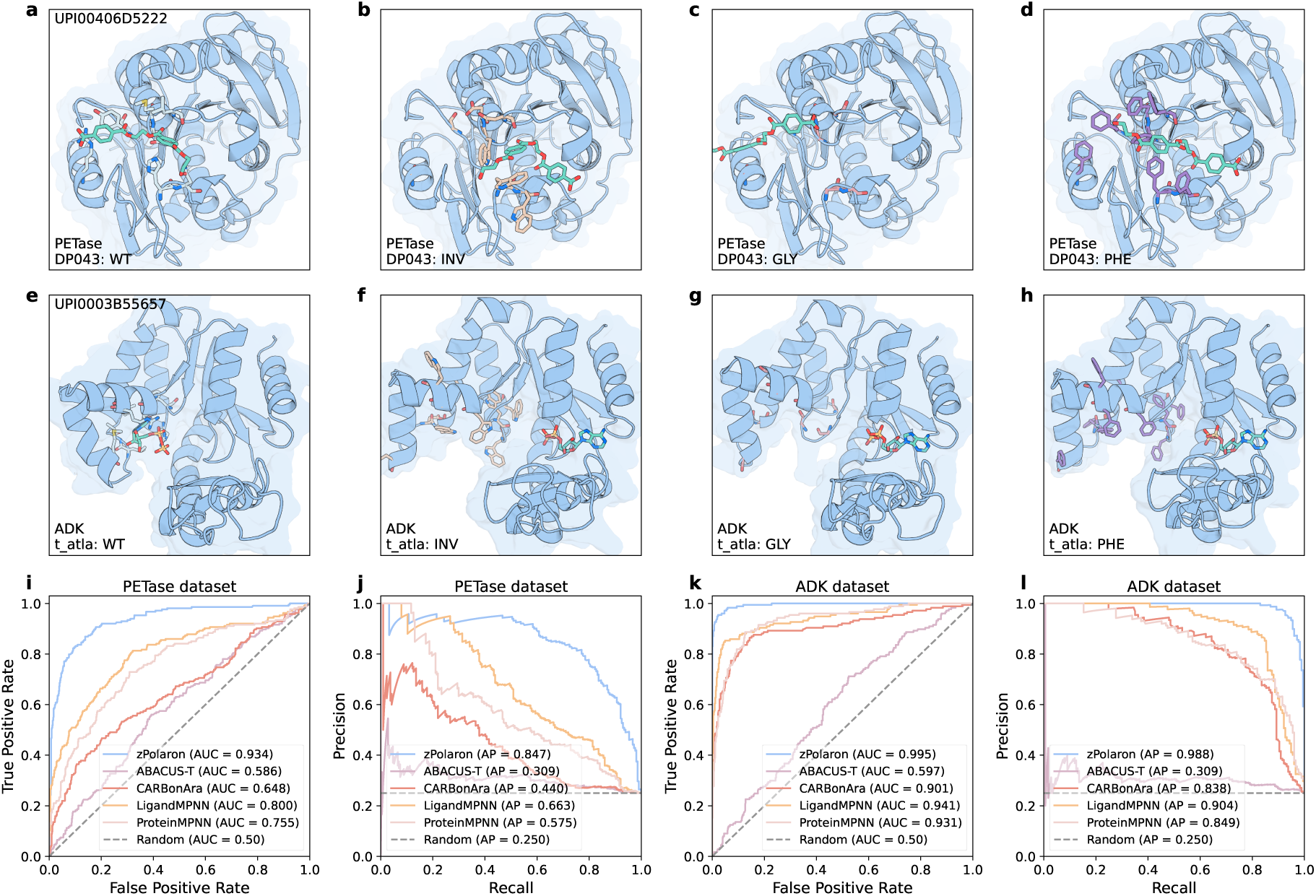
Performance on the disruptive binding-pocket mutation benchmark. (a–d) Representative AlphaFold 3-predicted structures of the PETase dataset after disruptive binding-pocket mutations: wild type (a), property inversion (b), side-chain removal (c), and hydrophobic packing (d). (e-h) Representative AlphaFold 3-predicted structures of the ADK dataset after disruptive binding-pocket mutations: wild type (e), property inversion (f), side-chain removal (g), and hydrophobic packing (h). (i, j) ROC (i) and precision–recall (j) curves of the five methods on the PETase dataset. (k, l) ROC (k) and precision–recall (l) curves of the five methods on the ADK dataset.

Notably, these three categories of disruptive mutations do not merely alter residues. They simultaneously perturb multiple aspects of protein function. GLY substitutions remove side-chain support and disrupt the local spatial geometry. PHE substitutions introduce severe steric hindrance, disrupting substrate access and binding. Property-inversion mutations directly disrupt the charge distribution and polarity network of the catalytic center. Therefore, these mutations concurrently involve constraints across multiple dimensions, including structural stability, non-covalent interactions, and evolutionary conservation. The advantage of zPolaron likely stems from its capability to jointly model this multilevel information. The graph neural network learns the 3D geometric relationships among residues, enabling the model to perceive spatial perturbations in the pocket structure. Meanwhile, the protein-ligand interaction graph explicitly encodes function-related constraints, including hydrogen bonding and hydrophobic interactions, allowing disruptions of the binding environment to be directly reflected in the model scores. Concurrently, the ESM2 embedding introduces evolutionary prior information from large-scale natural sequences. Because key catalytic residues in binding pockets are typically subject to strong selection over evolutionary timescales, they are much more conserved than ordinary residues. Consequently, these extreme mutations not only disrupt local structures and interaction networks but also deviate significantly from the sequence distribution shaped by the long-term evolution of the protein family. The integration of these three sources of information enables zPolaron to identify loss-of-function mutations more accurately, resulting in substantially better discrimination than the baseline methods.

### 3.5 Wet-lab validation confirms utility for enzyme discovery and redesign

The above results demonstrate that the zero-shot scoring strategy of zPolaron effectively captures changes in protein fitness and is highly sensitive to perturbations at functionally important residues. In practical protein engineering, however, the ultimate value of a computational model lies in its ability to leverage this information to discover enzymes with improved function or beneficial mutations. We therefore performed wet-lab experiments to evaluate the applicability of zPolaron in real-world enzyme engineering tasks.

To this end, we selected a two-step biocatalytic pathway for experimental validation. In this pathway, coniferyl alcohol is first oxidized to coniferaldehyde by CalA [71, 72], followed by the oxidation of coniferaldehyde to ferulic acid by CalB [71, 73]. All reactions were performed at 30 *^◦^*C and pH 9.0. The overall experimental workflow is illustrated in Fig. 5.

**Figure 5:**
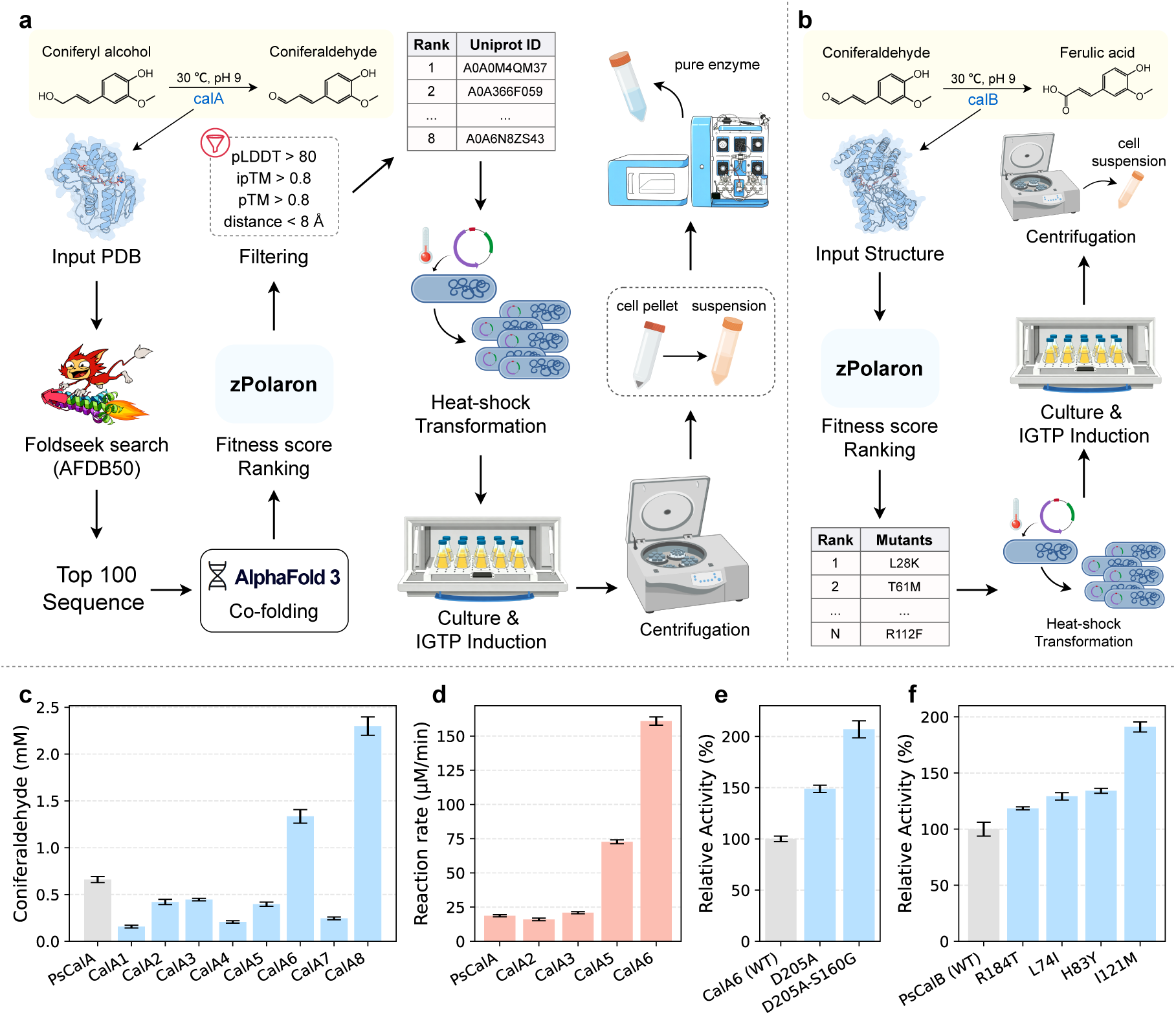
Workflow and experimental validation of enzyme discovery and mutation design. (a) Workflow for the enzyme discovery task. (b) Workflow for the mutation design task. (c) Coniferaldehyde production in whole-cell biotransformation catalyzed by PsCalA and the identified enzymes CalA1–CalA8. (d) Reaction rates of purified PsCalA and four selected CalA enzymes. (e) Relative activities of wildtype and mutant CalA6 measured by coniferaldehyde production in whole-cell biotransformation. (f) Relative activities of wild-type and mutant CalB measured by ferulic acid production in whole-cell biotransformation.

We first evaluated the ability of zPolaron to identify novel enzymes. The structure of the CalA-coniferyl alcohol-NAD^+^ complex was generated using AlphaFold 3 and used as the query for structural homology search against the AFDB50 database using Foldseek Server. The top 100 candidate proteins ranked by sequence similarity were retained, and their enzyme-substrate-cofactor complex structures were subsequently predicted using AlphaFold 3. These candidate complexes were then scored and ranked by zPolaron. To improve the reliability of candidate selection, additional structural quality filters were applied. Specifically, only models with an average confidence score greater than 80 and both pTM and ipTM values above 0.8 were retained. Furthermore, because the key step in the oxidation of coniferyl alcohol is hydride transfer from the substrate to the C4 atom of the nicotinamide ring in NAD^+^, we required the carbon atom bonded to the substrate hydroxyl group and the C4 atom of NAD^+^ to adopt a catalytically favorable geometry that approximates a near-attack conformation (NAC) (Fig. S4a). Since AlphaFold 3 does not always generate complex structures satisfying this criterion, additional structures were predicted using Boltz-2 [74] and included in the subsequent scoring and ranking procedure.

Candidate enzymes were selected for experimental validation based on the zPolaron scores together with the structural filtering criteria. The whole-cell biocatalytic activities of the selected candidates are shown in Fig. 5c. Reactions were carried out at 30 *^◦^*C and pH 9.0 for 30 min. CalA6 and CalA8 exhibited substantially higher whole-cell catalytic activities than the native CalA, whereas CalA2, CalA3, and CalA5 all retained more than 50% of the activity of CalA. Because heterologous expression levels may differ substantially among enzymes from different species, whole-cell activity alone cannot accurately reflect their intrinsic catalytic activities. Therefore, the five candidates with activities exceeding 50% of that of CalA were purified for in vitro enzymatic characterization [75]. Among them, CalA8 was predominantly expressed as inclusion bodies, preventing the purification of sufficient soluble protein for kinetic analysis.

Purified enzyme assays were performed under the same reaction conditions, and the results are shown in Fig. S5. Product concentrations increased approximately linearly over time for all enzymes. Therefore, the average reaction rate was calculated by linear regression of product concentration against reaction time, and the results are summarized in Fig. 5d. The average reaction rate of CalA was 18.68 *µ*M*/*min. CalA2 and CalA3 exhibited reaction rates of 15.99 and 20.97 *µ*M*/*min, respectively, comparable to that of CalA. In contrast, CalA5 reached 72.73 *µ*M*/*min, corresponding to a 3.89-fold improvement over CalA, while CalA6 achieved 160.97 *µ*M*/*min, representing an 8.62-fold increase. Overall, three candidate enzymes prioritized by zPolaron exhibited catalytic activities no lower than that of CalA, and two of them substantially outperformed the native enzyme.

Next, we evaluated the zero-shot prediction capability of zPolaron in enzyme mutation design. The overall workflow is shown in Fig. 5b. Directed mutagenesis was performed on both key enzymes involved in the two-step catalytic pathway. For CalA6, we first used zPolaron to score and rank all single-point mutations and experimentally evaluated the top-ranked candidates. The D205A mutant exhibited a catalytic activity 1.49-fold higher than that of the wild type. Based on this variant, saturation mutagenesis was subsequently performed at additional positions, followed by another round of ranking using zPolaron. Experimental measurements showed that the double mutant D205A-S160G further increased catalytic activity to 2.07-fold that of the wild type. Combined with the improvement obtained through enzyme discovery, the optimized variant CalA6-D205A-S160G ultimately achieved a catalytic rate 17.84 times higher than that of the native CalA.

We also optimized the enzyme responsible for the second step of the pathway, which catalyzes the oxidation of coniferaldehyde to ferulic acid. Candidate beneficial mutations predicted by zPolaron were constructed, heterologously expressed, and evaluated by whole-cell biocatalytic assays. Multiple mutants exhibited higher catalytic activity than the wild type. Among them, the I121M mutant showed the greatest improvement, producing 1.91 times more ferulic acid than the wild type under the same reaction conditions.

These experimental results demonstrate that zPolaron can not only identify novel enzymes with superior catalytic performance but also predict beneficial mutations that enhance enzyme activity. The model therefore shows strong potential for both enzyme discovery and enzyme optimization, validating its effectiveness as a zero-shot tool for enzyme design.

## 4 Discussion

### 4.1 Limitations of sequence recovery rate

Sequence recovery is the most widely used and intuitive evaluation metric for inverse folding models [15, 20, 69]. It measures the ability of a model to reconstruct the native amino acid identities within a given protein structure. However, this metric fundamentally reflects only the similarity between the designed sequence and the reference native sequence, and it is not directly equivalent to functional performance in protein engineering. Through these pre-trained models could be applied for stability and expression optimization [69, 68]. Our results further demonstrate that increased recovery rates do not necessarily lead to corresponding improvements in zero-shot fitness prediction or stability prediction. In other words, sequence recovery is more suitable for evaluating whether a model reproduces the native sequence distribution than for determining whether it has truly learned the constraints that govern protein function or fitness.

This limitation becomes particularly evident in different functional protein design scenarios. For tasks that aim to preserve the native fold, minimize sequence perturbation, and maintain family-level consensus constraints, a high sequence recovery rate is generally desirable. For example, in the design of extremozymes, when the starting protein is already naturally tolerant to high temperature, extreme pH, or organic solvents, higher sequence recovery generally indicates that the model is more likely to preserve key functional residues and environmental-related sequence distribution that have been retained through long-term natural selection [76, 43]. This helps maintain folding robustness and expression feasibility [15, 68]. Similarly, in conservative redesign, homologous protein repair, and stability optimization, high sequence recovery is typically more consistent with the design objective of introducing minimal sequence changes while preserving protein function.

In contrast, for applications that require exploration beyond the native sequence space, a lower sequence recovery rate is not necessarily a disadvantage and may even be essential [77]. For example, when the goal is to alter substrate specificity, remodel the binding pocket, introduce new catalytic activity, or explore novel functional combinations in distant sequence space, the designed sequences often need to deviate from the native consensus in order to establish new interaction patterns or adopt new conformational distributions [10, 11]. Therefore, sequence recovery should always be interpreted in the context of the specific design objective. It serves as a useful indicator of structural compatibility but should not be regarded as the sole criterion for evaluating the capability of an inverse folding model.

### 4.2 Evolutionary constraints beyond structural conservation

The objective of inverse folding is to learn the conditional probability distribution of a sequence given a protein backbone structure, *P* (sequence | structure), and to generate geometrically feasible and structurally compatible amino acid sequences from it [10]. Fundamentally, inverse folding models primarily focus on structural conservation, namely, whether the designed sequence can preserve the native three-dimensional fold of the target protein. Our results suggest that current state-of-the-art inverse folding models have largely achieved this objective. Whether using zPolaron, ProteinMPNN, LigandMPNN, or the other baseline methods, the designed sequences all produced highly confident structures when reconstructed by AlphaFold3, with consistently high TM-scores relative to the native crystal structures. In addition, the predicted secondary structures closely matched those of the native proteins. Together, these results indicate that existing inverse folding models are capable of preserving structural conservation.

However, the evolution of natural proteins is not simply a process of geometric optimization [38]. Instead, it results from long-term sampling of the fitness landscape under the combined effects of mutation, genetic drift, and natural selection [36]. Protein sequences are simultaneously shaped by functional activity, substrate recognition, folding kinetics, expression efficiency, and environmental adaptation [31, 32, 33, 78, 79]. Consequently, the structurally optimal sequence is not necessarily the functionally optimal one. For any given protein structure, many sequences may theoretically maintain the same folded state, while the function of these proteins are lost. The PSSM analysis in this study reflects this phenomenon. As the native PSSM score increases, the sequence recovery rate of zPolaron also increases monotonically, indicating that the model more accurately predicts highly conserved positions within natural protein families. However, the ΔPSSM analysis shows that the sequences generated by all inverse folding models exhibit an overall negative shift, meaning that the predicted residues generally have lower conservation scores than the corresponding native residues. This suggests that although these models successfully preserve the target structure, the designed sequences do not always follow the conservation patterns observed among natural homologs.

Sequence conservation may, in turn, be related to protein fitness [80]. Highly conserved residues in natural proteins often play essential roles in functional activity, and their conservation reflects fitness constraints accumulated through long-term natural selection. Nevertheless, protein fitness is not determined solely by the conservation of individual residues. Instead, it emerges from complex cooperative effects among multiple residues under specific functional selection pressures. Therefore, optimization based solely on structural information may produce thermodynamically reasonable sequences without fully capturing functional fitness. This interpretation is consistent with our experimental results. ProteinMPNN, as a representative structure-driven model, achieved strong performance in ΔΔ*G* prediction, indicating that it effectively captures constraints related to structural stability. However, when evaluated on fitness prediction tasks such as ProteinGym and DAOx, zPolaron consistently out-performed all baseline methods. The same advantage was observed in the identification of disruptive binding-pocket mutations and in subsequent experimental validation, where zPolaron demonstrated superior predictive performance and practical utility for protein engineering. These findings suggest that protein fitness is determined not only by structural stability but also by extensive function-related evolutionary constraints. The co-evolutionary relationships, family-level conservation patterns, and statistical regularities learned by protein language models from natural sequences provide complementary functional information beyond structural constraints [40, 41, 44]. Furthermore, protein–ligand interaction features supply local microenvironmental information that is directly related to protein function, enabling the model to balance structural compatibility with functional fitness.

Overall, current inverse folding models have become highly effective at preserving protein structure. However, structural conservation is not equivalent to sequence conservation, nor does it necessarily correspond to protein fitness. Future advances in inverse folding should therefore move beyond further improving sequence recovery alone. Rather than treating sequence recovery as a terminal objective, the next generation of inverse folding methods should fold evolutionary, functional, and molecular-interaction signals into the design criterion while preserving the structural scaffold. This redirects the field’s attention from structural compatibility toward functional fitness.

## 5 Conclusion

This study presents zPolaron, a heterogeneous graph neural network-based inverse folding model. Computational evaluation shows that zPolaron outperforms existing methods across multiple dimensions. In sequence recovery, zPolaron achieves the highest rate on an independent test set, with AlphaFold 3-validated TM-scores above 0.95. In evolutionary conservation analysis, zPolaron’s average ΔPSSM of *−*0.62 surpasses all baselines models, and its recovery rate increases monotonically with native PSSM scores. In zero-shot fitness prediction, zPolaron reaches Spearman correlations of *−*0.609 on Prote-inGym and maintains consistent values near *−*0.5 on the DAOx multi-substrate benchmark. Ablation experiments (Fig. S2) demonstrate that any two of the three information sources suffice to surpass all baselines. In disruptive mutation identification, zPolaron achieves AUC values of 0.934 and 0.995 on PETase and ADK, respectively, significantly outperforming all baselines. Wet-lab validation confirms the practical value of zPolaron. Through combined enzyme discovery and zero-shot mutation prediction, the optimized variant CalA6-D205A-S160G achieves a 17.84-fold improvement in catalytic activity over native CalA, and the CalB-I121M mutant shows a 1.91-fold increase over the wild type. In summary, structural conservation does not equate to protein fitness. These results indicate that integrating evolutionary and interaction information with structural constraints fills the missing puzzle of inverse folding, enhancing protein fitness prediction and moving sequence design beyond structural compatibility.

## 6 Data availability

The data that support this study are available from the corresponding authors upon request. The data and code are available at https://github.com/zelixirSH/zPolaron.

## 7 Acknowledgements

This work was supported by the National Key R&D Program of China (Grant No. 2023YFA0915500), the National Natural Science Foundation of China (Grant Nos. 12574225, 12275163, 32160217), the Taishan Scholars Program of Shandong Province (Grant No. tstp20240807, tsqn202312159), and the Yunnan Xingdian Talent Program (Youth Talent Program).

## 8 Contributions

L.Z. and W.L. designed and supervised the project. Z.W. designed the model and performed the computational and wet-lab experiments. Z.Z. performed the wet-lab experiments. Z.W., Z.Z., Y.Y., Y.Q., Y.M., S.W., N.H., L.Z., and W.L. analyzed data. Z.W., Z.Z., L.Z., and W.L. wrote the manuscript. Z.W., Z.Z, Y.Y., L.Z., and W.L. revised the manuscript.

