## Supplementary for "Filling the Missing Puzzle: Integrating Interaction and Evolutionary Information to Enhance Protein Fitness Prediction in Inverse Folding"

### 1 Features of zPolaron

#### 1.1 Node Features

Protein residue node features are formed by concatenating three components, with a total dimension of 667. Let the node feature of residue $r$ be $\mathbf{x}_r = [\mathbf{e}_r^{\mathrm{ESM}}; \mathbf{d}_r; \mathbf{o}_r] \in \mathbb{R}^{667}$, where each component is computed as:

$$\mathbf{e}_r^{\mathrm{ESM}} = \mathrm{ESM2}(\mathbf{S}_{\mathrm{masked}})[r] \in \mathbb{R}^{640} \tag{1}$$

$$\mathbf{d}_r = [\sin(\phi_r), \cos(\phi_r), \sin(\psi_r), \cos(\psi_r), \sin(\omega_r), \cos(\omega_r)] \in \mathbb{R}^6 \tag{2}$$

$$\mathbf{o}_r = \begin{cases} \text{one-hot}(21) & \text{if masked} \\ \text{one-hot}(a_r) & \text{otherwise} \end{cases} \in \mathbb{R}^{21} \tag{3}$$

where $\mathbf{S}_{\mathrm{masked}}$ is the protein sequence with residue $r$ replaced by a special mask token, $\phi_r, \psi_r, \omega_r$ are the three dihedral angles of residue $r$, and $a_r \in \{0, 1, \ldots, 19\}$ is the index of the 20 natural amino acid types for residue $r$ (not including the mask token).

Ligand atom node features employ a one-hot encoding representation of atom types, with a total dimension of 24. This encoding scheme classifies ligand atoms into 24 types, including nitrogen atoms of different hybridization states, carbon atoms of different hybridization states, oxygen atoms

*
†
‡
#These authors contributed equally to this work.

of different hybridization states, sulfur atoms of different hybridization states, hydrogen atoms, phosphorus atoms, halogen atoms, metal atoms, and other types of atoms. If the original type of an atom belongs to the halogen list, it is mapped to the halogen type; if it belongs to the metal list, it is mapped to the metal type; if it is not in the standard type list, it is mapped to the "other" type. This encoding scheme enables the model to distinguish atoms of different chemical properties within the ligand, thereby more accurately predicting their interactions with protein residues.

#### 1.2 Edge Features

The edges of the heterogeneous graph are divided into six types, each corresponding to distinct biological significance. Residue–residue peptide bond edges connect sequentially adjacent residues, encoding backbone continuity information. Residue–residue geometric edges connect spatially proximal residue pairs, without requiring sequential adjacency, used to capture long-range spatial interactions. Residue–residue interaction edges connect residue pairs with physicochemical interactions, with edge features encoding the type and strength of the interactions. Ligand–ligand covalent bond edges connect adjacent atoms within the ligand molecule, encoding the topological connectivity of the ligand. Ligand–residue geometric edges connect ligand atoms with spatially proximal protein residues, with edge features encoding their spatial relationships. Ligand–residue interaction edges connect ligand atoms and protein residues with non-covalent interactions, with edge features encoding the type and strength of the interactions.

##### 1.2.1 Interaction Edges

Protein–ligand interaction edge features encode 19 common types of non-covalent interactions, with each type occupying one dimension in the edge feature vector. Interaction determination follows the methods described in the work of Ferreira de Freitas and Schapira [1]. Interaction types include: five types related to hydrophobic interactions (aliphatic–aromatic, aromatic–aliphatic, aliphatic–aliphatic, carbon–halogen, sulfur–aromatic), two types related to hydrogen bonds (ligand donor–protein acceptor, protein donor–ligand acceptor), two types related to $\pi$–$\pi$ stacking (edge-to-face, face-to-face), two types related to weak hydrogen bonds (ligand donor–protein acceptor, protein donor–ligand acceptor), two types related to salt bridges (providing N atom and O atom, respectively), two types related to amide–aromatic stacking, two types related to cation–$\pi$ interactions, one type related to halogen bonds, and one type related to multipolar halogen interactions.

Protein internal interaction edge features contain 13 types of interactions, with each dimension of the edge feature vector representing the count of occurrences of that type of interaction between a given residue pair. Interaction types include: 3 types related to hydrophobic interactions (aliphatic–aromatic, aliphatic–aliphatic, sulfur–aromatic), 3 types related to hydrogen bonds (OH...O, NH...O, NH...N), 2 types related to $\pi$–$\pi$ stacking (edge-to-face, face-to-face), 2 types related to weak hydrogen bonds (aromatic and aliphatic carbon atoms as donors, respectively), 1 type related to salt bridges, and disulfide bonds (determined based on distance and angle criteria) as a non-covalent interaction.

The edge features are processed as follows: for each interaction type, geometric criteria (distance and angle thresholds) are used to determine the presence of the interaction. If present, the count value of the corresponding dimension is incremented by 1. Since a ligand atom may form the same type of interaction with multiple atoms within the same protein residue, the values in the edge feature vector are non-negative integer counts rather than simple 0/1 binary values. During model input preparation, interaction edge features associated with masked residues receive special treatment. Specifically, if an interaction edge connects to a masked residue, the feature vector of that edge is entirely set to a zero vector, while its original features are saved as training labels. This design prevents the model from seeing the original interaction information of masked residues during message passing, thereby avoiding information leakage. During training, the model is required to predict the original features of these masked edges as an auxiliary training task.

##### 1.2.2 Geometric Edges

**Residue–Residue Geometric Edge Features** This type of edge feature encodes the spatial relationship between two protein residues, with an edge feature vector dimension of 260. For the backbone atoms (C, O, N, CA) of residue $r_i$ and the backbone atoms (C, O, N, CA) of residue $r_j$, all sixteen pairs of inter-atomic distances are computed, yielding a $4 \times 4$ distance matrix. This distance vector is then expanded using radial basis functions:

$$\mathrm{RBF}(\mathbf{d}) = [\mathrm{RBF}(d_1), \ldots, \mathrm{RBF}(d_{16})] \in \mathbb{R}^{256} \tag{4}$$

where each distance value $d$ is expanded using sixteen radial basis functions:

$$\mathrm{RBF}(d) = [\exp(-\frac{(d-\mu_1)^2}{2\sigma^2}), \ldots, \exp(-\frac{(d-\mu_{16})^2}{2\sigma^2})] \tag{5}$$

with centers $\mu_k$ uniformly distributed between 0.0 and 20.0 Å and width $\sigma = 1.25$ Å. Next, directional projection features are computed. For each residue, a local coordinate frame is constructed using the N–CA–C atoms. The displacement vector from the virtual C$\beta$ atom [2, 3] of residue $r_j$ to that of residue $r_i$, $\Delta\mathbf{r}_{ij} = \mathbf{vCB}_j - \mathbf{vCB}_i$, is projected onto the three coordinate axes of residue $r_i$'s local frame, yielding a three-dimensional directional feature:

$$\mathbf{p}_{ij} = \frac{[\Delta\mathbf{r}_{ij} \cdot \mathbf{x}_i, \Delta\mathbf{r}_{ij} \cdot \mathbf{y}_i, \Delta\mathbf{r}_{ij} \cdot \mathbf{z}_i]}{\|\Delta\mathbf{r}_{ij}\|} \tag{6}$$

Finally, a distance sCalAr feature is computed by dividing the inter-residue distance by a cutoff threshold:

$$s_{ij} = \frac{\|\mathbf{vCB}_i - \mathbf{vCB}_j\|}{\text{distance\_threshold}} \tag{7}$$

Concatenating these three components yields the final geometric edge feature:

$$\mathbf{e}^{\text{geo}}_{r_i r_j} = [\mathrm{RBF}(\mathbf{d}_{ij}); \mathbf{p}_{ij}; s_{ij}] \in \mathbb{R}^{260} \tag{8}$$

**Ligand–Residue Geometric Edge Features** This type of edge feature encodes the spatial relationship between a ligand atom and a protein residue, with an edge feature vector dimension of 43. The distances from each ligand atom to the protein residue's backbone atoms and virtual C$\beta$ atom are computed, yielding a distance vector $\mathbf{d}_{lr} = [d_1, d_2, d_3, d_4, d_5] \in \mathbb{R}^5$. This distance vector is then expanded using radial basis functions:

$$\mathrm{RBF}(\mathbf{d}) = [\mathrm{RBF}(d_1), \ldots, \mathrm{RBF}(d_5)] \in \mathbb{R}^{40} \tag{9}$$

where each distance value $d$ is expanded using eight radial basis functions:

$$\mathrm{RBF}(d) = [\exp(-\frac{(d-\mu_1)^2}{2\sigma^2}), \ldots, \exp(-\frac{(d-\mu_8)^2}{2\sigma^2})] \tag{10}$$

with centers $\mu_k$ uniformly distributed between 0.0 and 8.0 Å and width $\sigma = 1.0$ Å. Next, directional projection features are computed similarly to the residue–residue geometric edge features. A local coordinate frame of the protein residue is first constructed, and the displacement vector from the residue virtual C$\beta$ atom to the ligand atom is projected onto the three coordinate axes:

$$\mathbf{p}_{lr} = \frac{[(\mathbf{p}_l - \mathbf{vCB}_r) \cdot \mathbf{x}_r, (\mathbf{p}_l - \mathbf{vCB}_r) \cdot \mathbf{y}_r, (\mathbf{p}_l - \mathbf{vCB}_r) \cdot \mathbf{z}_r]}{\|\mathbf{p}_l - \mathbf{vCB}_r\|} \tag{11}$$

Concatenating these two components yields the final geometric edge feature:

$$\mathbf{e}_{lr}^{\text{geo}} = [\text{RBF}(\mathbf{d}_{lr}); \mathbf{p}_{lr}] \in \mathbb{R}^{43} \tag{12}$$

##### 1.2.3 Peptide Bond Edges and Covalent Bond Edges

Peptide bond edges connect sequentially adjacent residues, encoding backbone continuity information. This type of edge has no additional feature vector; its mere existence indicates that two residues are sequentially adjacent and spatially connected by a peptide bond.

Ligand covalent bond edges connect adjacent atoms within the ligand molecule, encoding the topological connectivity of the ligand. This type of edge has no additional feature vector; its mere existence indicates that two atoms are connected by a covalent bond within the ligand molecule.

#### 2 Message Passing Neural Network

The core of zPolaron is a stacked heterogeneous graph neural network composed of multiple heterogeneous message-passing blocks connected in series [4]. Each block independently processes the six types of edges, aggregating messages from different neighbors to the central node through an attention mechanism, while dynamically updating edge features to capture higher-order interaction patterns.

Let the input node hidden state of the $l$-th block be $\mathbf{h}_v^{(l)}$ and the edge features be $\mathbf{e}_{uv}^{(l)}$. For each edge type $t$, the block first computes type-specific attention weights $\alpha_{uv}^{(t)}$ and messages $\mathbf{m}_{u\to v}^{(t)}$, then performs a weighted sum of messages across all types with a residual connection to the node's own state, and finally outputs the updated node hidden state $\mathbf{h}_v^{(l+1)}$ after layer normalization. The update formula for a heterogeneous message-passing block is:

$$\mathbf{h}_v^{(l+1)} = \text{LayerNorm}\left(\mathbf{h}_v^{(l)} + \sum_{t\in\mathcal{T}} \sum_{(u,v)\in E_t} \alpha_{uv}^{(t)} \cdot \text{MSG}^{(t)}(\mathbf{h}_u^{(l)}, \mathbf{h}_v^{(l)}, \mathbf{e}_{uv}^{(l)})\right) \quad (13)$$

where $\mathcal{T}$ denotes all edge types, $E_t$ is the set of edges of type $t$, and $\text{MSG}^{(t)}$ is the message function for type $t$. For the three internal residue edge types (peptide bond edges, geometric edges, interaction edges), the block uses graph attention convolution layers, attention cross-message-passing layers, and attention cross-message-passing layers, respectively. For intra-ligand edges, a graph attention convolution layer is used. For cross-type edges (ligand–residue geometric edges and interaction edges), attention cross-message-passing layers are used.

The attention cross-message-passing layer is the core component of message passing. This layer first maps edge features to the same dimensional space as node hidden states through linear projection, then concatenates them with source and target node hidden states as input to the message generation module. After message generation, multi-head attention assigns different importance weights to different neighbors, and finally the outputs of all heads are concatenated and fused through a linear transformation. The multi-head mechanism enables the model to capture neighbor importance from multiple different subspaces, improving the expressive power of message passing.

Edge features are updated together with node hidden states, reflecting the latest node representations at every layer. After processing by each message-passing block, the model uses the updated node hidden states to re-predict edge features:

$$\hat{\mathbf{e}}_{uv}^{(l+1)} = \text{MLP}_{\text{edge}}^{(t)}([\mathbf{h}_u^{(l+1)} \oplus \mathbf{h}_v^{(l+1)} \oplus \mathbf{e}_{uv}^{(l)}]) \quad (14)$$

where $\text{MLP}_{\text{edge}}^{(t)}$ is the edge feature update multi-layer perceptron for type $t$. This mechanism enables edge features to capture higher-order interaction patterns. The edge prediction task employs a zero-inflated Poisson loss function, composed of a binary classification loss and a Poisson negative log-likelihood loss.

The classifier module maps the node hidden states output by the final block to a 20-dimensional amino acid prediction space. Let the node hidden state output by the $L$-th block be $\mathbf{h}_v^{(L)} \in \mathbb{R}^{256}$. The classifier first maintains the hidden state at 256 dimensions through a set of linear layers with LayerNorm/ReLU/Dropout, then compresses it to a 128-dimensional feature space through the same components, and finally maps to the 20-dimensional space through a linear classification layer, yielding the predicted logits $\in \mathbb{R}^{20}$. During training, these logits are passed through a softmax activation to obtain predicted probability distributions for computing the classification loss.

### 3 Multi-Task Training Loss Strategy

zPolaron is trained end-to-end in a multi-task learning framework, with the total loss defined as the sum of the node classification loss and a dynamically weighted, sparsity-regularized edge prediction loss:

$$\mathcal{L}_{\text{total}} = \mathcal{L}_{\text{node}} + \mathcal{L}^{*}_{\text{edge}} \tag{15}$$

where $\mathcal{L}^{*}_{\text{edge}}$ incorporates both dynamic weight scheduling and sparsity regularization:

$$\mathcal{L}^{*}_{\text{edge}} = w_{\text{edge}}(e)\, \mathcal{L}_{\text{edge}} + \lambda_{\text{sparse}}\, \mathcal{L}_{\text{sparse}} \tag{16}$$

with $w_{\text{edge}}(e) = \lambda_{\text{edge}} \min\bigl(1,\ (e+1)/T_{\text{warmup}}\bigr)$, $\lambda_{\text{edge}} = 8$, $T_{\text{warmup}} = 10$, and $\lambda_{\text{sparse}} = 10^{-5}$.

#### 3.1 Node Classification Loss

The node classification loss trains the model to predict amino acid identity at each residue position and integrates three design components that jointly address class imbalance, hard-sample learning, and overconfidence.

**Label smoothing with distribution awareness.** Standard label smoothing converts hard one-hot targets to soft targets, mitigating overconfidence. zPolaron extends this with a distribution-aware mechanism that adjusts the smoothing strength per class according to its frequency in the training set: high-frequency classes receive stronger smoothing, low-frequency classes weaker smoothing. The smoothed target reduces the probability of the true class slightly below 1 and raises the probabilities of the remaining classes slightly above 0, without altering the relative ranking.

**Focal loss for hard-sample emphasis.** Building on the smoothed cross-entropy, a focal term $(1 - p_{i,y_i})^{\gamma}$ is introduced to down-weight well-classified (easy) samples, redirecting training focus toward hard samples. Here $p_{i,y_i}$ denotes the model's predicted probability for the true amino acid $y_i$ at position $i$, and $\gamma \geq 0$ is the focal parameter. When $\gamma = 0$, the loss reduces to standard smoothed cross-entropy; as $\gamma$ increases, the contribution of easy samples is progressively suppressed.

**Entropy regularization.** A penalty on the entropy of the predicted distribution $H(\mathbf{p}_i) = -\sum_{j=0}^{19} p_{ij} \log p_{ij}$ is subtracted from the loss, encouraging flatter (higher-entropy) output distributions and further guarding against overconfident predictions.

The node classification loss additionally applies class-specific weights $w_{y_i}$ to compensate for amino acid frequency imbalance. The final form is:

$$\mathcal{L}_{\text{node}} = \frac{\sum_{i=1}^{N} w_{y_i}\, (1 - p_{i,y_i})^{\gamma} \Bigl[-\sum_{j=0}^{19} t_{ij} \log p_{ij}\Bigr]}{\sum_{i=1}^{N} w_{y_i}} \; - \; \lambda_{\text{conf}}\, \frac{1}{N} \sum_{i=1}^{N} H(\mathbf{p}_i) \tag{17}$$

#### 3.2 Edge Prediction Loss

The edge prediction loss trains the model to estimate interaction-type counts between residue pairs and between residues and ligand atoms. Because the vast majority of residue or atom pairs lack a given interaction type (count = 0), the count data exhibit severe zero-inflation—zeros far outnumber what an ordinary Poisson distribution would predict. We therefore employ a zero-inflated Poisson (ZIP) loss, which models the data as a mixture of two latent processes: a binary component that governs whether a count is a structural zero (i.e., necessarily zero), and a Poisson component that generates the actual count for non-zero cases. This mixture formulation substantially improves the fit to zero-inflated interaction data.

The ZIP loss has the form:

$$\mathcal{L}_{\text{edge}} = \alpha \cdot \text{BCE}(\pi, z) + (1 - \alpha) \cdot \mathcal{L}_{\text{Poisson}} \tag{18}$$

where $\pi$ (output via a sigmoid) is the predicted probability that the sample belongs to the structural-zero category, $z \in \{0, 1\}$ is the binary indicator of whether the true count is zero, and $\mathcal{L}_{\text{Poisson}}$ is the Poisson negative log-likelihood evaluated on the non-zero samples.

### 4 Datasets

#### 4.1 Dataset for Sequence Recovery (PDB2023-2025)

Table S1: Test set of zPolaron.

| | | | | | | | | | | | |
|---|---|---|---|---|---|---|---|---|---|---|---|
| 7qp5 | 7r1m | 7tlw | 7uba | 7uka | 7uni | 7x54 | 7xlf | 7xn2 | 7xps | 7xra | 7xrb |
| 7xrx | 7y9c | 7yf2 | 7yf3 | 7yf4 | 7yib | 7ype | 7yug | 8a2g | 8a2n | 8abu | 8agy |
| 8akh | 8an0 | 8avq | 8b61 | 8bau | 8bc5 | 8brp | 8btj | 8bux | 8c7p | 8cjw | 8cp2 |
| 8cpn | 8d00 | 8d2z | 8dot | 8dq2 | 8e5f | 8e5g | 8ebb | 8ehn | 8eil | 8eip | 8evk |
| 8ew8 | 8f3k | 8fai | 8fts | 8fzb | 8g1n | 8g52 | 8g64 | 8gbk | 8gdl | 8gmh | 8grs |
| 8gth | 8gyw | 8gze | 8h3z | 8h4q | 8haw | 8hja | 8hkr | 8hw5 | 8hwi | 8ifu | 8ii8 |
| 8iqd | 8ixw | 8j2f | 8j5h | 8j8k | 8jbi | 8ji2 | 8jjn | 8jkz | 8jsc | 8jsk | 8jul |
| 8jvf | 8jx4 | 8jx6 | 8k2m | 8k66 | 8kbj | 8khg | 8on5 | 8ou1 | 8p0s | 8pak | 8paq |
| 8pu1 | 8pu4 | 8q28 | 8q8o | 8qag | 8qta | 8r34 | 8rcw | 8rdd | 8rv3 | 8ryd | 8s1j |
| 8s3m | 8s3u | 8s3w | 8smq | 8sx0 | 8sx2 | 8syj | 8szb | 8t8k | 8t8y | 8tl8 | 8tn6 |
| 8tuk | 8twq | 8u01 | 8uus | 8ux2 | 8v3m | 8v3o | 8v4w | 8vc8 | 8vdx | 8vej | 8whl |
| 8wnw | 8wtl | 8x3z | 8x4m | 8x4n | 8x5e | 8xgq | 8xif | 8xij | 8xkd | 8xma | 8xpf |
| 8xsn | 8xua | 8yf3 | 8yjp | 8yk9 | 8yv5 | 8yzy | 8z4q | 8zia | 8zic | 8zij | 8zlf |
| 8zxr | 9azz | 9b8e | 9ba3 | 9bj5 | 9c5w | 9c9n | 9ca3 | 9cd2 | 9cg2 | 9cjk | 9d5x |
| 9dcs | 9de4 | 9dq2 | 9dyq | 9e0s | 9e50 | 9eav | 9ebm | 9efd | 9em9 | 9emt | 9evl |
| 9f5q | 9f9v | 9fiz | 9fw5 | 9fyi | 9gcz | 9gmj | 9gn5 | 9grz | 9h62 | 9hle | 9i2l |
| 9i4z | 9iab | 9ik4 | 9ink | 9iqd | 9iqu | 9ir1 | 9irx | 9iur | 9j0v | 9j15 | 9j33 |
| 9j7d | 9j7v | 9jbh | 9jn8 | 9jqf | 9jtz | 9k3o | 9k92 | 9kbo | 9kct | 9kkb | 9ko3 |
| 9kqj | 9kqv | 9kw9 | 9l34 | 9l93 | 9lb6 | 9ljg | 9lp2 | 9m7f | 9mno | 9mrb | 9mrg |
| 9n1s | 9n4g | 9ntn | 9nto | 9omi | 9ow2 | 9pd0 | 9q2y | 9qvr | 9r2l | 9r2o | 9sp7 |
| 9srg | 9u8h | 9u9y | 9uba | 9uu1 | 9vww | 9vzi | 9w7g | 9y02 | | | |

#### 4.2 PDB entries for structure re-prediction using AlphaFold 3

Table S2: PDB entries for structure re-prediction using AlphaFold 3.

| | | | | | | | | | | | |
|---|---|---|---|---|---|---|---|---|---|---|---|
| 7qp5 | 7r1m | 7tlw | 7xlf | 7xn2 | 7xps | 7y9c | 7yf2 | 7yf3 | 7yib | 7yug | 8akh |
| 8bau | 8c7p | 8cp2 | 8d00 | 8d2z | 8e5g | 8ebb | 8ehn | 8evk | 8ew8 | 8f3k | 8fts |
| 8g52 | 8g64 | 8gdl | 8gyw | 8gze | 8ifu | 8j8k | 8jsc | 8jsk | 8k2m | 8khg | 8ou1 |
| 8pak | 8paq | 8q28 | 8qta | 8rcw | 8rdd | 8s1j | 8s3m | 8s3u | 8s3w | 8sx0 | 8t8k |
| 8t8y | 8tn6 | 8vc8 | 8vdx | 8x4m | 8xif | 8xkd | 8yjp | 8yv5 | 8z4q | 8zic | 8zlf |
| 8zxr | 9azz | 9cd2 | 9cg2 | 9de4 | 9dq2 | 9dyq | 9ebm | 9emt | 9evl | 9fiz | 9gmj |
| 9i2l | 9iab | 9ink | 9ir1 | 9j0v | 9j15 | 9j33 | 9j7v | 9jbh | 9k3o | 9kct | 9l34 |
| 9lb6 | 9n1s | 9n4g | 9q2y | 9qvr | 9sp7 | 9srg | 9u8h | 9vzi | | | |

AlphaFold 3 is sensitive to system complexity, so we pre-filtered structures to keep the downstream analysis trustworthy. We therefore applied the following criteria to the PDB2023–2025 dataset: (1) the number of residues per protein chain is constrained to between 30 and 1,000—the lower bound excludes short peptide fragments that lack a well-defined folded structure, while the upper bound avoids systematic biases introduced by computational resource limitations and elevated prediction uncertainty for very long chains; (2) each PDB entry contains no more than five protein chains, to control the combinatorial complexity of multi-chain systems and reduce the risk of prediction errors at chain–chain interfaces; (3) no more than three consecutive residues are missing within a single chain, preventing structural gaps from propagating local conformational discontinuities to neighboring regions. At the prediction stage, multi-chain complexes are further decomposed into individual single-chain inputs, eliminating inter-chain dependencies that could confound structure prediction.

#### 4.3 Disruptive Binding-Pocket Mutation Dataset

Two datasets were used for the disruptive binding-pocket mutation analysis. The ADK dataset and the PETase dataset. Following the protocol described by Masters *et al.* [5], we introduced three types of disruptive mutations into residues located within the binding pocket. The mutation strategy was identical to that used in our previous work.

Specifically, AlphaFold 3 was first used to predict the co-folded structure of each enzyme-substrate complex. The minimum distance between every substrate atom and every protein atom was then calculated. A residue was defined as a binding-pocket residue if the shortest distance between any of its atoms and any substrate atom was within 4 Å. All identified pocket residues were subsequently subjected to disruptive mutations according to the following rules:

(1) Side-chain erasure (GLY mutation): The critical residues are mutated to glycine to eliminate side-chain steric support, thereby simulating the structural collapse of the pocket and the loss of non-covalent anchoring points.

(2) Hydrophobic packing (PHE mutation): The residues are mutated to bulky phenylalanine, inducing physical crowding and steric hindrance within the active cavity to obstruct the substrate-binding channel.

(3) Property inversion (INV mutation): The residues are mutated into amino acids with completely opposite physicochemical or charge properties, thoroughly disrupting the original catalytic polar network.

Table S3: Assignment for mutating to dissimilar binding site residues.

| Wild-Type | Mutation | Wild-Type | Mutation |
|---|---|---|---|
| TYR | GLY | GLY | TRP |
| TRP | GLY | PRO | TRP |
| LYS | GLY | HIS | TRP |
| ARG | GLY | ASP | TRP |
| VAL | ASP | GLU | TRP |
| LEU | ASP | CYS | TRP |
| ILE | ASP | ASN | TRP |
| MET | ASP | GLN | TRP |
| PHE | ASP | THR | TRP |
| ALA | TRP | SER | TRP |

#### 4.4 P450 dataset for fine-tuning

To further evaluate the adaptability of zPolaron to a specific enzyme family, we constructed a dataset based on cytochrome P450 (CYP) enzymes. All enzyme sequences and their corresponding reaction information were obtained from the P450Rdb v2.0 database [6]. For each enzyme in the database, the protein sequence, heme cofactor (HEM), and corresponding substrate were provided as inputs to AlphaFold 3 to predict the enzyme-HEM-substrate ternary complex structure. Only samples with successfully generated complex structures (i.e., complete CIF files) were retained for subsequent analyses.

The amino acid sequence of each sample was extracted from the protein sequence provided by P450Rdb. To avoid high sequence homology between the training and test sets, all sequences were clustered using MMseqs2 (easy-linclust) with a minimum sequence identity threshold of 0.4 [7]. After clustering, all samples within the same cluster were treated as an indivisible data unit, ensuring that no sequences sharing more than 40% identity were distributed across different datasets.

Because some P450 families were substantially larger than others, the training data could otherwise become heavily biased toward a few dominant families. To alleviate this imbalance, at most 20 samples were randomly retained from each cluster. The clusters were then randomly divided into training, validation, and test sets at an approximate ratio of 8:1:1. All samples belonging to the same cluster were assigned to the same subset, thereby preventing performance overestimation caused by the leakage of highly homologous sequences.

The resulting training, validation, and test sets all consisted of AlphaFold 3-predicted enzyme-HEM-substrate complex structures. Because the entire dataset was constructed from predicted rather than experimentally determined structures, it was used to evaluate both the applicability of zPolaron to AlphaFold 3-generated structures and its transfer learning capability for a specific enzyme family. During fine-tuning, only the output classification layer was updated using the training set, while the backbone network remained frozen, allowing us to assess whether the model could rapidly learn P450 family-specific sequence preferences while preserving its general structural representations.

### 5 Ablation study

To evaluate the contribution of each core component of zPolaron to sequence design and function prediction, we designed three ablation experiments that remove three key sources of information from the model.

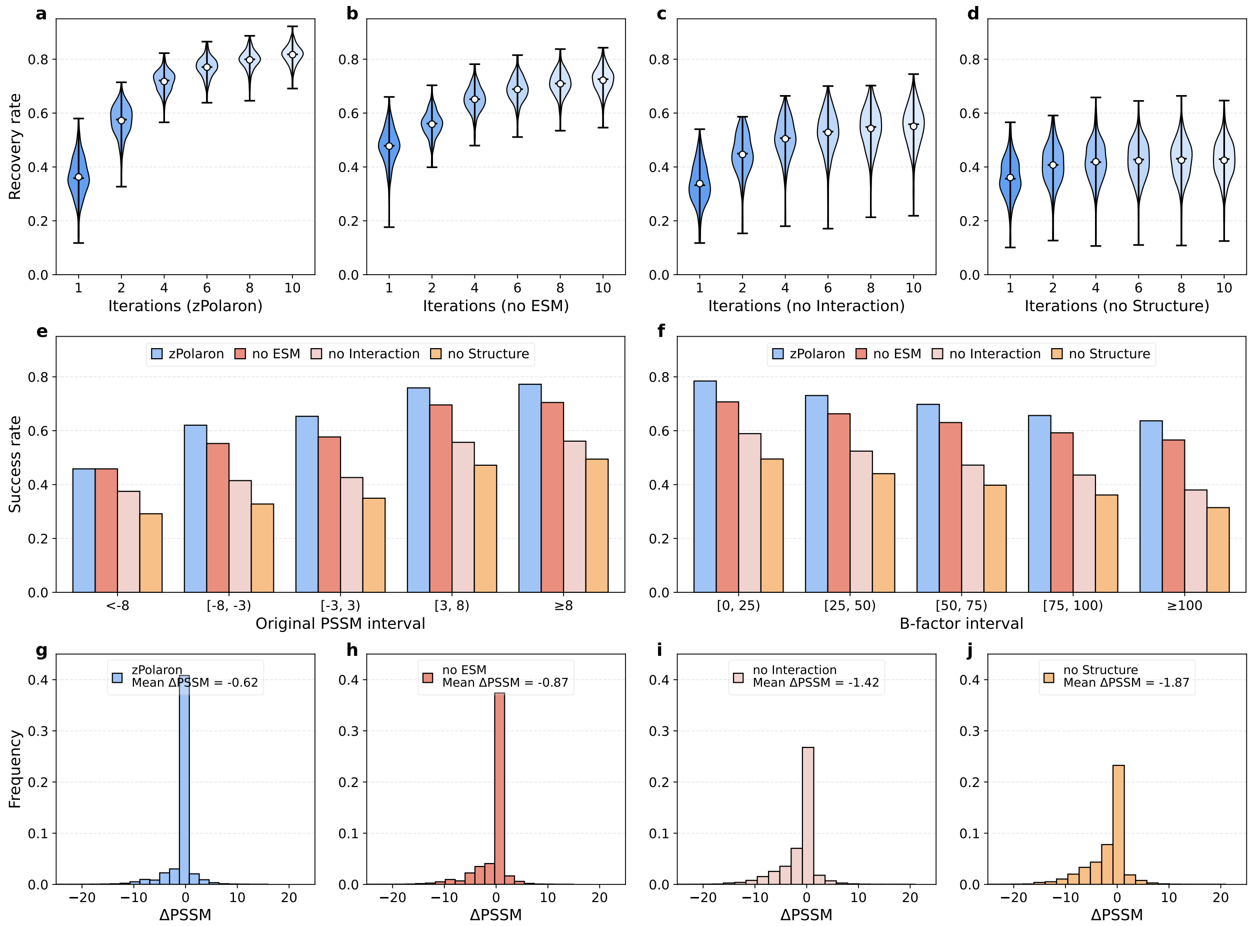


Figure S1: Sequence recovery and residue-level performance of the ablation models.(a-d) Sequence recovery over successive design iterations for the full zPolaron model (a), zPolaron without ESM embeddings (b), zPolaron without interaction features (c), and zPolaron without backbone structural features (d). (e) Residue recovery rate across different native residue PSSM intervals. (f) Residue recovery rate across different B-factor intervals. (g-j) Distributions of ΔPSSM, defined as the difference between the PSSM score of the predicted residue and that of the corresponding native residue.

(1) Removal of ESM2 evolutionary embeddings (no ESM). The ESM2 pre-trained language model embeddings in the residue node features are all set to zero, retaining only the dihedral angle geometric features and amino acid identity one-hot encoding. This experiment tests whether evolutionary prior information provides incremental contributions beyond structural geometry encoding.

(2) Removal of interaction edge features (no Interaction). All edge attributes of inter-residue and residue-ligand interaction edges are set to zero, and the auxiliary training task for edge prediction is also removed. The model relies solely on covalent bond edges (peptide bonds, ligand chemical bonds) and geometric edges for message passing. This experiment tests whether explicit

non-covalent interaction encoding (hydrogen bonds, hydrophobic interactions, $\pi$–$\pi$ stacking, etc.) provides chemical constraints that cannot be replaced by geometric distance information.

(3) Removal of backbone structure features (no Structure). The backbone dihedral angles (sin/cos, 6 dimensions) and amino acid identity one-hot encoding (21 dimensions) in the residue node features are all set to zero, retaining only the ESM2 embeddings. This experiment tests the necessity of explicit structural geometry features—that is, whether the implicit structural awareness of the language model can replace explicit geometric encoding.

The training data and hyperparameters of the ablated models remain consistent with those of the full model, and during inference, the same confidence-guided iterative strategy as the full model is adopted.

The sequence recovery rates of the ablation models decreased in the order of the full model > zPolaron (no ESM) > zPolaron (no Interaction) > zPolaron (no Structure). Notably, zPolaron (no ESM) achieved a slightly higher initial recovery rate than all other models in the first iteration, indicating that the initial ESM2 embeddings, when all residues are masked, primarily encode sequence-length information and may introduce priors that are inconsistent with structural constraints. The iterative refinement mechanism effectively corrected this issue by progressively incorporating confident contextual information, allowing the full model to surpass zPolaron (no ESM) from the second iteration onward. In contrast, zPolaron (no Structure) exhibited the lowest recovery rate, reaching only 42.49% after ten iterations, with little improvement throughout the iterative process. This result suggests that the effectiveness of the confidence-guided strategy is highly dependent on the model's underlying prediction capability.

For all ablation variants, sequence recovery increased with increasing PSSM scores. The full model consistently achieved the highest recovery rates, indicating that each module contributes positively to the recognition of evolutionary signals. This trend was further quantified by the $\Delta$PSSM analysis. The full model achieved an average $\Delta$PSSM of $-0.62$, substantially outperforming all three ablation variants.

Regarding robustness to structural uncertainty, zPolaron (no Structure) exhibited the most severe performance degradation, demonstrating that explicit geometric encoding is the most critical component of the inverse folding framework and the primary source of its robustness against structural noise. Removing either ESM2-derived features or interaction features also led to noticeable performance declines, suggesting that these complementary sources of information provide additional constraints when structural information becomes increasingly uncertain.

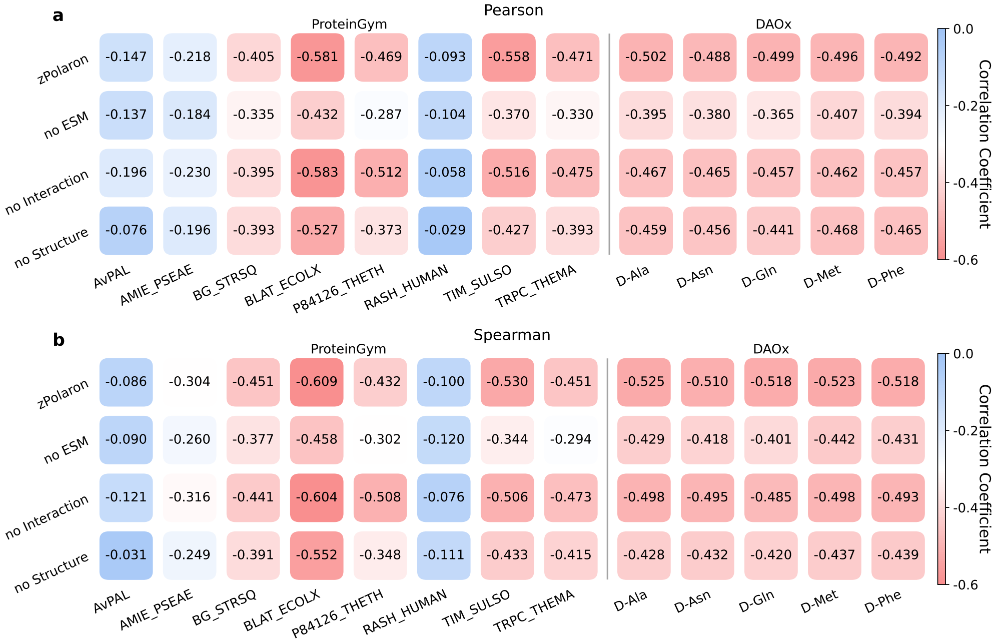


Figure S2: Correlation analysis of zero-shot scoring in the ablation study. (a) Pearson correlations between zero-shot scores and protein fitness values on protein fitness benchmark datasets. (b) Spearman correlations between zero-shot scores and protein fitness values on protein fitness benchmark datasets.

### 6 Generalization to the Cytochrome P450 enzyme family

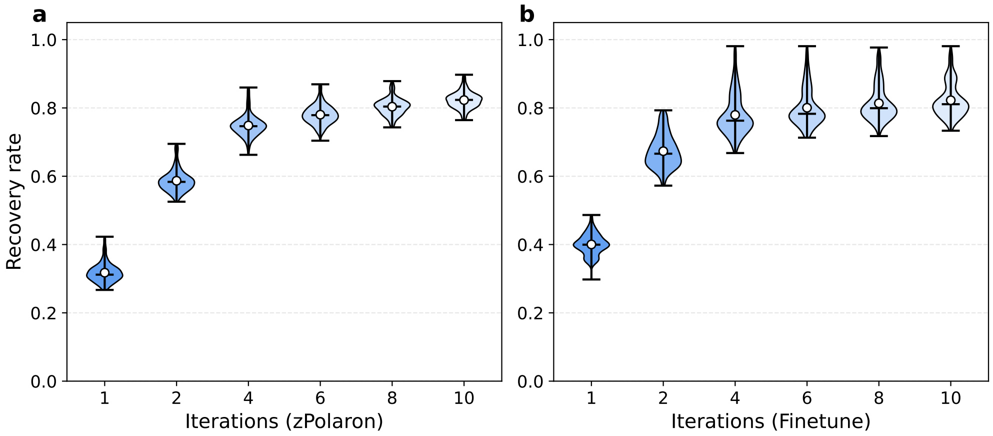


Figure S3: Sequence recovery on the P450 enzyme dataset. (a) Sequence recovery of zPolaron on the P450 enzyme dataset without fine-tuning. (b) Sequence recovery of fine-tuned zPolaron on the P450 enzyme dataset.

To further evaluate the scalability of zPolaron for specific protein families, we performed an additional study using the cytochrome P450 (CYP) enzyme family. Cytochrome P450 represents one of the most extensively studied enzyme families in protein engineering because of its broad substrate scope and diverse catalytic functions, making it a major target for enzyme engineering and functional design. However, high-resolution experimental structures are unavailable for many CYP homologs. Consequently, protein engineering studies often rely on structure prediction methods such as AlphaFold 3 to obtain three-dimensional structures. In this study, we therefore used AlphaFold 3-predicted CYP structures as model inputs to examine whether zPolaron can be directly applied to predicted structures rather than being limited to experimentally determined ones.

We first applied the pretrained zPolaron model directly to the CYP dataset and evaluated sequence recovery across different confidence-guided iterative design rounds. Despite the absence of any family-specific training, the model achieved competitive sequence recovery on AlphaFold 3-predicted structures, and the recovery rate increased progressively with successive iterations. These results indicate that the structural representations learned by zPolaron generalize well to predicted protein structures and that the confidence-guided iterative strategy remains effective in this setting.

Next, we performed lightweight fine-tuning by updating only the output classification layer while keeping the backbone network fixed. Compared with the pretrained model, the fine-tuned model consistently achieved higher sequence recovery at every iteration. This result suggests that the learned latent representations are highly transferable across protein families, whereas family-specific sequence preferences can be efficiently captured by adapting only a small number of task-specific parameters.

Overall, these results demonstrate that zPolaron is applicable not only to inverse folding based on experimentally determined PDB structures but also to AlphaFold 3-predicted structures. Moreover, the model can be rapidly adapted to specific protein families using a limited amount of family-specific data, highlighting its generalizability and scalability for practical protein engineering applications.

### 7 t-SNE Dimensionality Reduction Results of zPolaron Embeddings

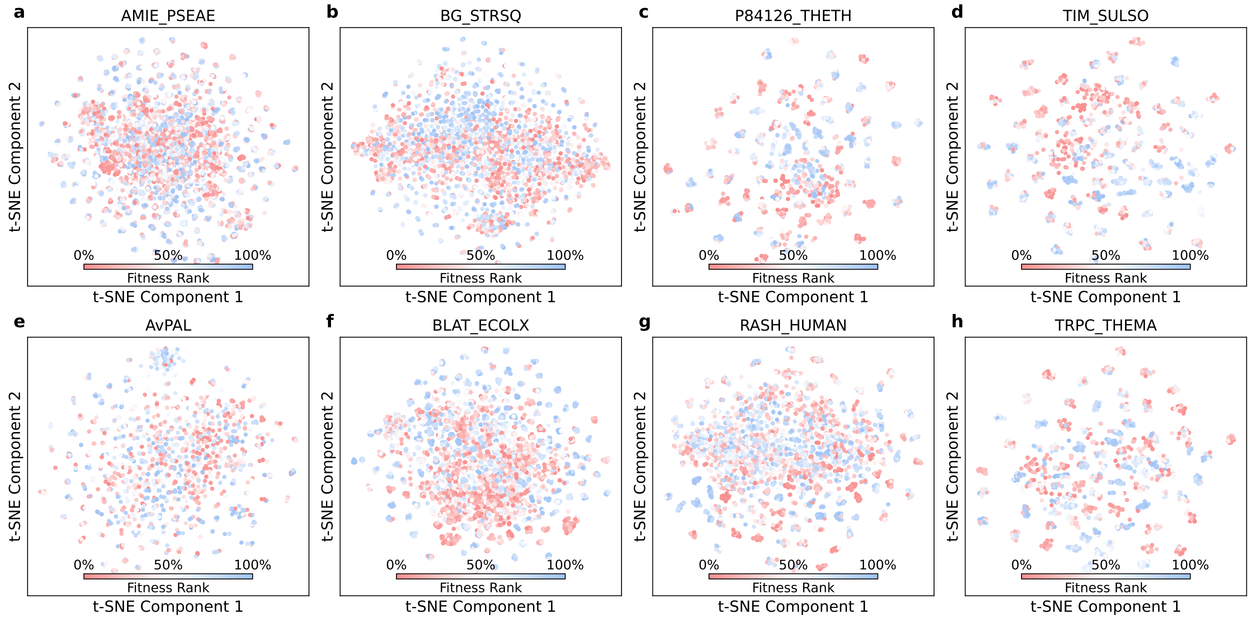


Figure S4: Results of t-SNE dimensionality reduction of zPolaron embeddings across 8 different targets in ProteinGym.

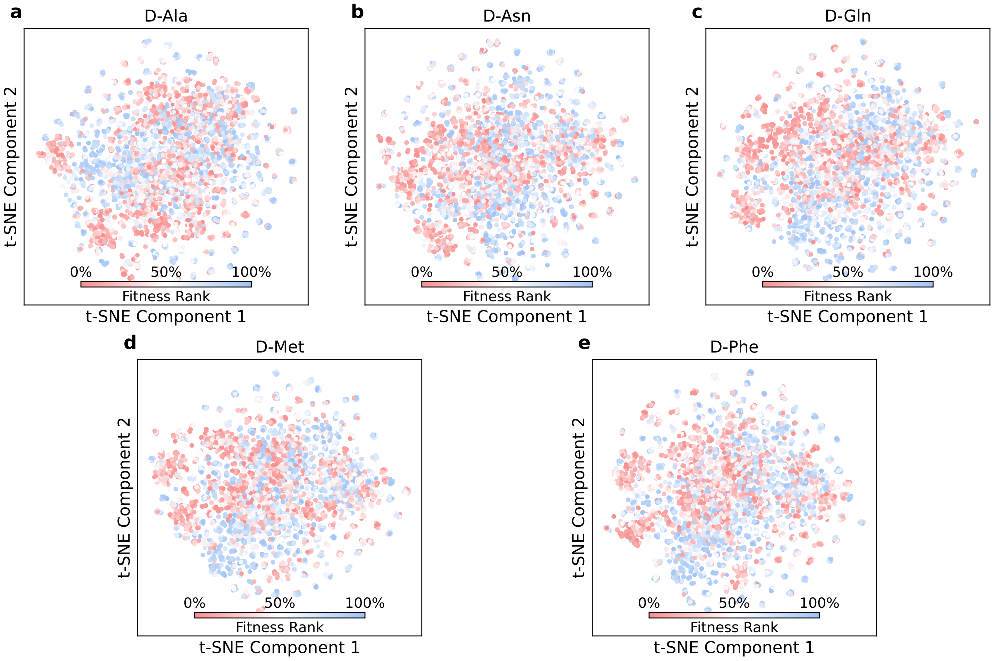


Figure S5: Results of t-SNE dimensionality reduction of zPolaron embeddings across 5 different substrates of DAOx.

### 8 Materials and methods for wet-lab experiment

#### 8.1 SDS-PAGE gel image

The images below show representative SDS-PAGE gels of samples collected at various stages of purification using a protein purification system. From left to right, the lanes correspond to: molecular weight marker, whole cell lysate, lysate pellet, lysate supernatant, flow-through, contaminant proteins, purified enzyme, and empty vector control.

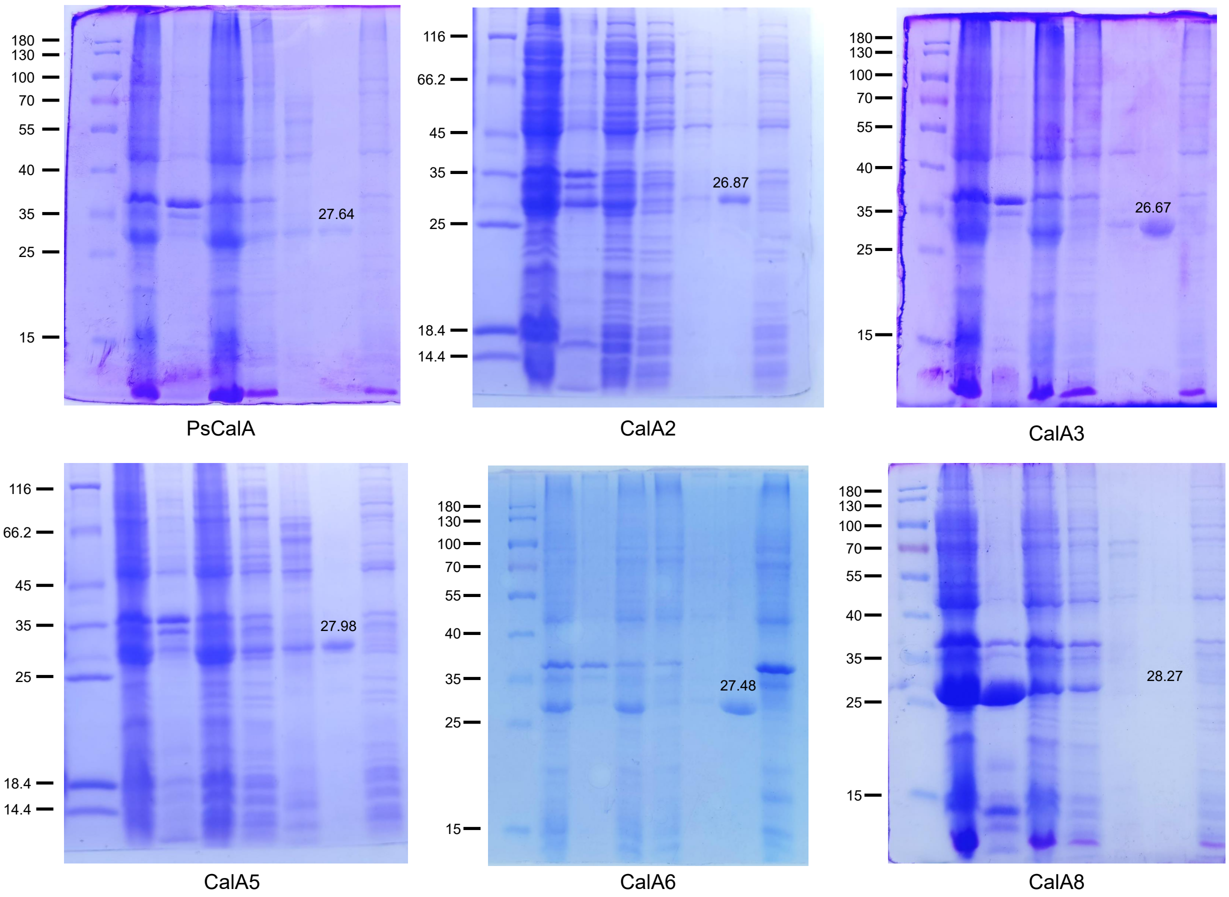


Figure S6: SDS-PAGE gel image of purified PsCalA, CalA2, CalA3, CalA5, CalA6, and CalA8.

#### 8.2 High-performance liquid chromatography (HPLC) conditions and standard curve

The HPLC system used was a Shimadzu LC-20AD equipped with a Hypersil$^{TM}$ BDS C18 column (250 mm length, 4.6 mm inner diameter, 4 µM particle size). The mobile phase consisted of 0.1% formic acid in water (mobile phase A) and acetonitrile (mobile phase B), with an isocratic ratio of A:B = 2:8. The standard solutions for the calibration curve were prepared using analytical-grade standards, following the same preparation procedure as that used for the experimental samples.

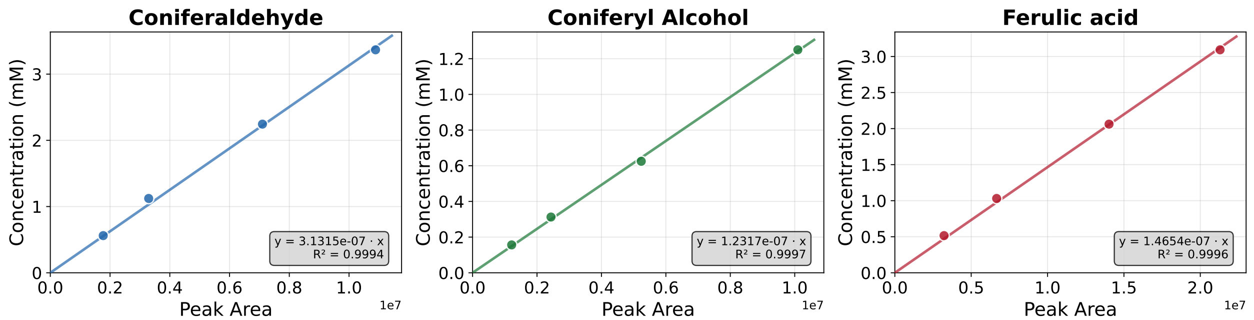


Figure S7: HPLC Standard curve.

##### 8.3 Amino acid sequence and nucleotide sequence

Table S4: Enzyme information.

| Name | UniprotID | Species Origin |
|---|---|---|
| PsCalA | P0DMP5 | Pseudomonas sp. (strain HR199 / DSM 7063) |
| CalA1 | A0A0M4QM37 | Pseudonocardia sp. HH130629-09 |
| CalA2 | A0A366F059 | Roseiarcus fermentans |
| CalA3 | A0A2T7UV07 | Pararhodobacter aggregans |
| CalA4 | A0A7Y8KE50 | Pseudomonas edaphica |
| CalA5 | A0A523M326 | Gammaproteobacteria bacterium |
| CalA6 | A0A2C9T3J5 | Mycobacterium sp. shizuoka-1 |
| CalA7 | A0A527X7A8 | Mesorhizobium sp |
| CalA8 | A0A6N8ZS43 | Gammaproteobacteria bacterium |
| PsCalB | A0A2A2GIR2 | Paracoccus salipaludis |

**PsCalA** Amino acid sequence:

MQLTNKKIVVTGVSSGIGAETARVLRSHGATVIGVDRNMPSLTLDAFVQADLSHPEGIDKAISQLPEKIDGLCNIAGVPGTADPQLVANVNYLGLKYLTEAVLSRIQPGGSIVNVSSVLGAEWPARLQLHKELGSVVGFSEGQAWLKQNPVAPEFCYQYFKEALIVWSQVQAQEWFMRTSVRMNCIAPGPVFTPILNEFVTMLGQERTQADAHRIKRPAYADEVAAVIAFMCAEESRWINGINIPVDGGLASTYV

**PsCalA** Nucleotide sequence:

ATGCAACTCACTAACAAAAAGATTGTTGTTACCGGTGTTTCTTCTGGTATTGGTGCGGAAACCGCGCGTGTTCTGCGTTCTCACGGTGCGACCGTTATCGGTGTTGACCGTAACATGCCGTCTCTGACCCTGGACGCGTTTGTTCAGGCTGACCTGTCTCACCCAGAGGGTATCGACAAAGCGATCTCTCAGCTGCCGGAAAAGATCGACGGTCTGTGCAACATCGCGGGTGTTCCGGGTACCGCGGACCCGCAGCTGGTTGCGAACGTTAACTACCTGGGTCTGAAATACCTGACCGAAGCGGTTCTCTCTCGTATCCAGCCGGGTGGTTCTATCGTTAATGTGTCTTCTGTGCTGGGCGCTGAATGGCCAGCGCGTCTGCAACTGCACAAAGAACTGGGTTCCGTTGTTGGTTTCTCTGAAGGTCAGGCGTGGCTGAAGCAGAACCCGGTTGCGCCGGAATTTTGCTACCAGTACTTCAAAGAAGCGCTGATCGTTTGGTCTCAGGTGCAGGCGCAGGAATGGTTCATGCGTACCTCTGTTCGTATGAACTGCATCGCGCCAGGTCCAGTTTTCACCCCGATCCTGAACGAGTTCGTGACTATGCTGGGTCAGGAACGTACCCAAGCGGACGCGCATCGTATCAAACGTCCGGCGTATGCGGACGAAGTTGCGGCGGTTATCGCGTTCATGTGCGCGGAAGAATCTCGTTGGATCAACGGTATCAACATTCCGGTTGACGGTGGTCTGGCGTCTACCTACGTT

**CalA1** Amino acid sequence:

MTTDPTGVRRFVVTGAGSGIGAALTDRLVTSGHEVVALDRAVSGVPDGAEPLVCDRTDPAAVEAAVAAVRATGPLHGVAAVAGVPGSVPPPVVHAVNVVGLRRVVGALAPLLAPGSAVVLLSSMAGYRGTATPDELDRYLTLGDDDLQAALEATALDGPGAY

QLSKQVVHALAADLAATLHPRGVRAVSVSPGPVETPILADFRATMPTLDEASRLVGRNARAEEVAAVVEFVLSPAASWLNGIDLRLDGGLTALRARAVR

**CalA1** Nucleotide sequence:

ATGACCACCGACCCAACCGGCGTTCGCCGTTTTGTGGTCACCGGTGCAGGTAGCGGTATTGGCGCGGCGCTCACCGACCGTCTGGTCACCAGCGGTCATGAAGTGGTGGCGCTTGACCGCGCGGTTAGCGGCGTGCCGGACGGCGCTGAACCGCTGGTCTGTGACCGTACCGATCCTGCAGCGGTTGAAGCGGCAGTTGCTGCAGTTCGCGCAACCGGCCCGCTTCACGGCGTTGCGGCCGTTGCAGGCGTTCCGGGCAGTGTTCCGCCGCCGGTGGTTCATGCGGTGAACGTGGTCGGTCTGCGCCGCGTGGTCGGCGCGCTTGCGCCGCTGCTGGCGCCGGGCAGTGCGGTGGTGCTGCTGTCCTCTATGGCCGGTTATCGCGGCACGGCAACGCCGGATGAACTTGACCGCTATCTCACCCTGGGCGATGATGATCTTCAGGCGGCGCTTGAAGCCACGGCGCTTGACGGCCCGGGCGCTTATCAGCTGTCTAAACAGGTGGTTCATGCGCTGGCGGCAGATCTTGCGGCAACGCTGCATCCGCGCGGCGTGCGTGCGGTGTCCGTTAGTCCAGGCCCGGTTGAAACGCCGATTTTGGCTGATTTTCGCGCCACCATGCCGACCCTTGATGAAGCGAGCCGCCTGGTGGGCCGTAATGCCCGTGCTGAAGAAGTTGCTGCGGTGGTGGAGTTTGTGCTGTCCCCTGCAGCAAGCTGGCTGAACGGTATTGATCTCCGTCTTGATGGCGGCTTGACCGCGCTGCGCGCCCGCGCGGTGCGC

**CalA2** Amino acid sequence:

MLFGKTIVVTGIASGIGARIGQLALALGADVVGLDVNAPASPTGEFVRLDIASAAEIEAVVKTLPARVDALCNVAGVSGVSGAQKTLAINFYGLRALTEAVAPRLREGGAIVNVASIAGYGWRANLERAKAFVAAPGFPDIAALLVEHNVPDGEGYPLSKELLLLWTQQAAHQPLFRGRGLRVNAVSPGPVTTPIFPEFRKLFGDPRVDDDIARVGRAGSAADIAPAALFLCSDGARWINGANLPVDGGLEASINATALGF

**CalA2** Nucleotide sequence:

ATGTTGTTTGGTAAAACCATTGTGGTCACCGGTATTGCGAGCGGTATTGGCGCGCGTATTGGGCAGCTGGCGCTGGCGCTGGGCGCAGATGTGGTCGGTCTTGATGTTAATGCGCCTGCATCGCCAACCGGTGAATTTGTCCGTTTGGATATTGCCAGTGCGGCAGAAATTGAAGCGGTGGTGAAAACCCTTCCTGCACGCGTTGATGCGCTCTGTAACGTTGCGGGCGTTAGTGGCGTTTCAGGCGCGCAAAAAACGTTGGCGATTAACTTTTACGGCCTGCGGGCGTTGACCGAAGCGGTTGCGCCGCGTTTGCGTGAAGGCGGCGCGATCGTTAACGTCGCGTCGATTGCCGGTTATGGCTGGCGCGCGAATCTTGAACGCGCAAAAGCGTTTGTTGCCGCGCCAGGTTTCCCGGATATTGCCGCGCTGCTGGTTGAGCATAACGTGCCGGATGGCGAAGGTTATCCGCTGAGTAAAGAACTGCTGCTGCTCTGGACGCAGCAGGCGGCGCATCAGCCGCTGTTTCGCGGCCGCGGCCTTCGCGTGAACGCGGTTTCTCCAGGCCCGGTAACCACGCCAATTTTCCCGGAATTCCGCAAACTGTTTGGCGATCCGCGCGTTGATGATGATATTGCCCGCGTTGGCCGTGCAGGTAGTGCGGCAGATATTGCGCCTGCGGCGCTGTTTCTGTGTTCTGATGGCGCGCGCTGGATTAACGGCGCGAATCTCCCGGTTGATGGCGGTCTTGAAGCGTCGATTAACGCCACCGCGCTGGGTTTT

**CalA3** Amino acid sequence:

MRVIVTGAASGIGRAIAERLLADGHQVVALDLRPVGLPGAECHALDQGDPASIDRVLSAIAGPLDGLVNSAGLPPRPGNEAQLLRVNLLGLKHLTEAVLDRLTPGGAVVTIASRAGHAWRENLTQARAALALGWHDDVAGFCAREGIDALRAYALSKEALILWTRQSAAPLIARDLRMNALCPAAVETPLLEDFARLQSAESIARNAARTARRGRVADIAPVAAFLLSPESGWIKGEALALDGGLSAMAECEALGL

**CalA3** Nucleotide sequence:

ATGCGCGTGATCGTCACCGGCGCGGCGAGCGGTATTGGCCGCGCGATTGCCGAACGCTTGCTGGCCGACGGTCATCAGGTTGTGGCGCTGGATCTCCGCCCGGTTGGCTTGCCGGGCGCGGAATGTCATGCGCTTGATCAGGGCGATCCGGCGAGTATTGACCGCGTGTTGAGCGCCATTGCCGGGCCGCTTGATGGCCTGGTGAACAGCGCCGGTTTGCCGCCGCGTCCGGGCAATGAAGCGCAGCTGCTGCGCGTGAACCTGTTGGGCCTTAAACATCTGACCGAAGCGGTGCTGGACCGCCTGACGCCGGGCGGCGCGGTGGTGACCATCGCCTCCCGCGCCGGTCATGCGTGGCGCGAAAACCTCACCCAGGCCCGCGCGGCGCTGGCGCTTGGCTGGCATGATGATGTGGCCGGTTTCTGCGCCCGTGAAGGCATTGATGCGTTGCGCGCTTATGCGCTGTCAAAAGAAGCGCTGATTCTCTGGACCCGCCAGTCGGCCGCGCCGTTGATTGCCCGCGATCTCCGCATGAACGCCCTCTGCCCGGCGGCGGTTGAAACGCCGCTGCTGGAAGATTTTGCCCGTTTGCAGAGTGCAGAATCGATTGCCCGTAATGCGGCACGTACCGCGCGTCGCGGCCGCGTGGCGGATATTGCGCCGGTTGCGGCATTTCTGCTGTCGCCGGAATCCGGCTGGATTAAGGGCGAAGCGCTGGCGCTTGATGGCGGCTTGTCCGCGATGGCTGAATGTGAAGCGCTGGGCTTG

**CalA4** Amino acid sequence:

LPARVHGLCNIAGVPGTAPVHAVATVNYLGLRHLTQSLLPRMPAGGSIVNVASVLGAQWPQRLELHKALAAIPSFEAGQQWLAANPVEQGSCYQYFKEALIVWSFQQSQGWFRDHSVRINCVAPGPVFTPILGDFVSMLGPQRVAEDSQRMTRPALADEVAAV

IAFLCSDAARWVNGVNLPVDGGLAATYV

**CalA4** Nucleotide sequence:

TTGCCTGCCCGCGTTCACGGCTTGTGCAATATTGCCGGCGTGCCGGGCACCGCGCCGGTTCATGCGGTCGCGACGGT
GAACTATCTCGGCTTGCGCCATTTGACGCAGAGTTTGTTGCCGCGTATGCCTGCAGGCGGCTCGATCGTGAACGTCGCGA
GCGTGCTTGGCGCGCAGTGGCCGCAACGTCTTGAATTGCATAAAGCGCTTGCGGCAATTCCGTCGTTTGAAGCCGGGCA
GCAGTGGCTGGCGGCAAATCCGGTTGAGCAGGGCAGCTGCTATCAGTATTTTAAAGAAGCGTTGATCGTCTGGAGTTTC
CAGCAGTCGCAGGGCTGGTTTCGCGATCATAGCGTGCGCATTAACTGCGTTGCGCCAGGCCCGGTATTTACGCCGATTCT
TGGTGATTTTGTGTCGATGCTTGGCCCGCAGCGCGTGGCGGAAGATTCGCAGCGCATGACCCGCCCGGCGCTGGCTGAT
GAAGTTGCGGCGGTGATTGCGTTTCTCTGTAGTGATGCCGCACGCTGGGTGAACGGCGTGAATCTGCCGGTTGATGGCG
GTTTGGCGGCAACTTATGTT

**CalA5** Amino acid sequence:

MKVIVVTGGATGMGAATVTRLVAADCEVHVLDVNPPEAIEVRFHRCDLGSPAAIDATVANLPAAIDGLINVAGIALAEPPE
QTLAVNFLGLRHLTEACLPRTREGGSVVNVASTAGWDWRKREPVVRELLATPDFPAGLDWLRANEAEWIDNPYKFSKQCAAA
YTYRAAGLALNGRVRVNCVNPGAVETQLTPAFRELVGPALYDWGVEQIGRQGCPDDVAEVIEFLSIGQCGWLNGVEIPVDGGY
LAGLTGGWINPDESPKD

**CalA5** Nucleotide sequence:

ATGAAAGTGATTGTGGTCACCGGCGGCGCAACCGGTATGGGCGCGGCAACGGTCACCCGTTTGGTTGCTGCAGACT
GTGAAGTTCATGTGCTGGATGTTAATCCGCCAGAAGCGATTGAAGTCCGTTTTCACCGCTGTGATCTTGGCAGTCCTGCG
GCGATTGATGCCACCGTCGCAAATCTTCCTGCGGCGATTGATGGCCTGATTAACGTTGCGGGTATTGCGCTTGCCGAACC
GCCAGAACAAACGCTGGCGGTGAACTTTTTGGGCTTGCGGCATCTGACCGAAGCCTGTCTGCCGCGTACCCGTGAAGGC
GGCAGTGTGGTGAACGTCGCGTCAACCGCAGGCTGGGACTGGCGCAAACGCGAACCGGTGGTGCGCGAATTGCTGGCAA
CGCCAGATTTCCCTGCAGGTCTTGACTGGTTGCGCGCGAACGAAGCCGAATGGATTGATAACCCGTATAAATTCTCCAAA
CAGTGTGCGGCGGCTTATACCTACCGCGCGGCAGGTCTGGCGCTGAACGGCCGCGTGCGCGTGAACTGCGTTAACCCGG
GCGCGGTTGAAACGCAACTGACGCCAGCATTTCGCGAACTGGTTGGCCCGGCGCTTTATGACTGGGGCGTTGAGCAGAT
TGGCCGTCAGGGCTGCCCGGATGATGTGGCTGAAGTGATTGAATTTCTGTCCATCGGTCAGTGCGGCTGGCTTAACGGC
GTTGAAATTCCGGTTGATGGCGGCTATCTCGCCGGCTTGACCGGCGGCTGGATTAATCCGGACGAATCGCCGAAAGAT

**CalA6** Amino acid sequence:

MASFSGRRYVVTGAASGIGHAVAEQLLALGAEVTSLDRNTPTAPVQAHIEVDLANPRSIDAALEQLNGDYDGLLNVAGIPGT
AASELVFAVNSLAVRHLTEAFFDRLVPGGSVTIVSSTAGFGWPERLGVIRDLLATDTFEEGAAWYKANPQQGNAYNFSKEVTTV
YALSMGLAMAQMGFRINAVLPGPVQTPILVDFEDTMGKDTLDGLKELLGRHAEPEDIAELVLFLASDAARWVNGQAIAVDGGIS
GAVASGVVPAPEI

**CalA6** Nucleotide sequence:

ATGGCCAGTTTTAGCGGCCGCCGTTATGTGGTCACCGGCGCGGCAAGCGGTATTGGTCATGCGGTTGCTGAACAGTT
GCTTGCGCTGGGCGCAGAAGTCACCAGTCTTGACCGTAATACGCCAACCGCGCCGGTTCAGGCGCATATTGAAGTTGATT
TGGCAAATCCGCGCTCAATTGATGCGGCGCTTGAGCAGCTTAACGGCGATTATGACGGCCTGCTTAACGTTGCGGGTATT
CCGGGCACTGCAGCATCTGAACTGGTTTTTGCGGTGAACTCTCTTGCCGTTCGGCATCTGACCGAAGCGTTTTTTGACCG
CCTGGTTCCGGGCGGCAGCGTCACCATCGTCTCCTCCACCGCAGGTTTTGGCTGGCCGGAACGCCTTGGCGTGATCCGCG
ATTTGCTGGCGACCGATACCTTTGAAGAAGGCGCGGCCTGGTATAAAGCCAATCCGCAGCAGGGCAATGCCTATAACTTC
TCTAAAGAAGTCACCACCGTTTATGCGCTGAGTATGGGCCTGGCGATGGCGCAGATGGGTTTTCGTATTAACGCCGTGCT
GCCAGGCCCGGTACAAACGCCGATTTTGGTGGATTTTGAAGACACCATGGGTAAAGACACCCTTGATGGCTTGAAAGAAC
TGCTGGGCCGTCATGCTGAACCGGAAGATATTGCCGAACTGGTGCTGTTTCTGGCCTCTGATGCCGCACGCTGGGTGAAC
GGTCAGGCGATTGCCGTTGATGGCGGTATTAGCGGCGCGGTTGCGAGTGGCGTTGTTCCTGCACCGGAAATC

**CalA7** Amino acid sequence:

MLFGKTILITGVASGIGARTAELAGQLGADVIGVDMREPADHQGTFVKADISSRAGVDDLVRQLPQRLDALCNVAGLSGNT
GAVPTLAVNFYGLRALSEALAPKLREGGAIVNVASIAGFGWRANLNRAASMVSVEGFPDVARVIAQHAVKNEEGYPVSKELLLL
WTFRAAHQEQFKSRGIRVNAVSPGPVETP

**CalA7** Nucleotide sequence:

ATGTTGTTTGGTAAAACCATTCTGATTACCGGCGTTGCGAGCGGTATTGGCGCACGTACCGCAGAACTTGCAGGTCA

GTTGGGCGCGGATGTGATTGGCGTTGATATGCGCGAACCTGCAGATCATCAGGGCACGTTTGTTAAAGCCGATATCAGC
AGCCGCGCGGGCGTTGATGATCTGGTCCGCCAGTTGCCGCAACGTCTTGATGCGCTCTGTAACGTTGCGGGTTTGAGCG
GCAATACCGGCGCGGTACCGACGCTGGCGGTGAACTTCTACGGCCTTCGCGCGCTGAGCGAAGCGCTTGCGCCGAAACT
GCGCGAAGGCGGCGCGATCGTGAACGTCGCGTCGATTGCCGGTTTTGGCTGGCGCGCGAACCTCAACCGCGCGGCGAGT
ATGGTGAGCGTTGAAGGTTTCCCGGATGTGGCGCGCGTGATTGCGCAGCATGCGGTGAAAAACGAAGAAGGCTATCCGG
TGAGTAAAGAACTGCTGCTGCTCTGGACCTTCCGCGCGGCGCATCAGGAGCAGTTTAAAAGCCGCGGTATTCGCGTGAAT
GCGGTTTCGCCAGGCCCGGTTGAAACGCCG

**CalA8** Amino acid sequence:

MDLAAKRIVVTGAASGIGKESARMLRERGATVIGLDRKQPANGLVDQFIRVDLTEFESIGRAVNAIEGKVDGLCNIAGVPPT
ASIATVVTVNFIALRYLTERMLDKLADGASIINMASLAGVGWPDSIEQVKRFIAEADFNNADSLCEELGIDEERSYFFAKEVLRVW
TMQNWNTWRQRGIRVNAISPGPVETPILADFLETLGERAEEDMGVMGRAGTVEEIAPIVVFLCSDDSVWINGTNIAADGGMQA
HVMRQVHGF

**CalA8** Nucleotide sequence:

ATGGATCTTGCCGCAAAACGCATTGTGGTCACCGGCGCGGCAAGCGGTATTGGTAAAGAATCCGCCCGTATGTTGCG
CGAACGCGGCGCAACGGTTATTGGTCTTGACCGTAAACAGCCGGCAAATGGCCTGGTTGATCAGTTTATTCGCGTTGATC
TGACCGAATTTGAATCTATTGGCCGCGCGGTAAATGCCATTGAAGGCAAAGTTGATGGCCTGTGTAATATTGCCGGCGTT
CCGCCAACGGCAAGTATTGCCACCGTCGTCACCGTGAACTTTATCGCCCTGCGGTATCTGACCGAACGCATGCTCGATAA
ACTGGCTGATGGCGCGAGCATCATCAATATGGCCTCGCTTGCGGGCGTTGGCTGGCCGGATTCTATTGAACAGGTGAAA
CGCTTTATTGCCGAAGCTGATTTTAATAATGCCGATAGCTTGTGTGAAGAACTCGGCATTGATGAAGAACGCAGCTATTT
CTTCGCCAAAGAAGTGCTTCGCGTCTGGACCATGCAGAACTGGAATACCTGGCGCCAGCGCGGTATTCGCGTGAACGCCA
TTTCGCCAGGCCCGGTTGAAACGCCGATTCTGGCTGATTTTCTCGAAACCTTGGGCGAACGCGCTGAAGAAGATATGGGC
GTTATGGGCCGTGCAGGTACGGTTGAAGAAATTGCGCCGATTGTGGTGTTTCTGTGTTCTGATGATTCCGTCTGGATTAA
CGGCACCAATATTGCCGCTGATGGCGGTATGCAGGCGCATGTTATGCGTCAGGTTCACGGTTTC

**PsCalB** Amino acid sequence:

MSQTTASLDPDHPALALAFRHLSDAAGPRPDVTDRRKSLAALARALGRHEDGLIAAVDADFGGRTAAETRLLELLPLYDQI
AHARRHLRGWMRRRRVAGSRFLWPARAFYEYQPLGVVGVIGAWNYPVLLTLGPMVDALAAGNRVLVKPSELAPRTAEALAAL
VAEAFPPGQVAVVTGDAALARALTRLPLDHLVFTGSTRVGREVMRAAADNLTPVTLELGGKSPAILHDSADLGRAVPRLMTGK
LLNAGQTCVAPDYALVPPSRMAEFEAEARRAVAALFPNGLNGDYTRIISDRHLERLKGLVAEAREGGARVVELAPVPAGPNGRL
MPPMLVFDAPEDSALMREEIFGPILPVIPAPTLDDALAFVNARPRPLALYCFAEDGARQDEVLARTMSGGVTFNDCMFHVGQLN
LPFGGVGASGTGEYHGFDGFARFSKKRPVMVQPRLASTGLARPPWGARRRLIGAMLAFARRAARR

**PsCalB** Nucleotide sequence:

ATGTCGCAAACCACCGCAAGTCTTGATCCTGATCATCCCGCGCTTGCGCTGGCATTTCGCCATCTGTCTGATGCCGC
AGGCCCGCGGCCGGATGTGACCGACCGCCGCAAATCGCTTGCCGCGCTTGCCCGCGCGCTTGGCCGCCATGAAGATGGCC
TGATTGCCGCCGTTGATGCCGATTTTGGCGGCCGTACGGCGGCAGAAACCCGCCTGCTGGAACTGCTGCCGCTTTATGAC
CAAATTGCCCATGCCCGCCGCCATCTTCGCGGCTGGATGCGCCGCCGCCGCGTTGCGGGTAGCCGTTTTCTCTGGCCGGC
GCGCGCGTTTTATGAATATCAGCCGCTGGGCGTGGTCGGCGTGATTGGCGCGTGGAACTATCCGGTGCTGCTGACCCTG
GGGCCGATGGTTGATGCGCTGGCGGCAGGTAACAGGGTGCTGGTTAAACCGTCCGAACTGGCGCCGCGTACCGCAGAAG
CGCTGGCGGCGCTGGTGGCAGAAGCATTCCCGCCAGGGCAGGTGGCGGTGGTGACCGGCGATGCGGCGCTTGCGCGCGC
GCTAACCCGCCTGCCGCTTGATCATCTGGTCTTTACCGGCTCCACCCGCGTTGGCCGCGAAGTGATGCGCGCGGCGGCAG
ATAACCTCACGCCGGTCACCCTGGAACTGGGCGGTAAATCCCCGGCGATTCTTCATGATAGTGCCGATCTGGGCCGCGCG
GTTCCGCGCCTGATGACCGGCAAATTGCTGAACGCCGGGCAGACCTGCGTTGCGCCGGATTATGCGTTGGTTCCGCCGTC
ACGTATGGCTGAATTTGAAGCCGAAGCCCGCCGCGCGGTTGCGGCGTTGTTCCCGAACGGCTTGAACGGCGATTACACCC
GCATTATCAGCGACCGCCATTTGGAACGCCTGAAAGGCTTGGTGGCTGAAGCCCGTGAAGGCGGCGCGCGCGTGGTGGA
ACTTGCGCCGGTTCCTGCAGGGCCGAACGGCCGCCTTATGCCGCCGATGCTGGTGTTTGATGCGCCGGAAGATTCGGCGC
TTATGCGCGAAGAAATTTTTGGCCCGATTCTGCCGGTGATCCCTGCGCCAACGCTGGATGATGCGCTGGCGTTTGTGAAT
GCCCGCCCGCGCCCGCTGGCGCTTTACTGCTTTGCCGAAGATGGCGCGCGTCAGGATGAAGTGCTTGCCCGCACCATGAG
CGGCGGCGTGACCTTTAACGACTGCATGTTCCACGTTGGGCAGCTGAATCTGCCGTTTGGCGGCGTTGGCGCGTCCGGTA
CCGGCGAATATCACGGCTTTGACGGTTTTGCCCGTTTTAGCAAAAAACGCCCGGTGATGGTTCAGCCGCGCCTGGCGTCA

ACCGGTCTGGCCCGGCCGCCGTGGGGCGCGCCGCCGCTTGATTGGCGCGATGCTGGCGTTTGCCCGCCGCGCGGCAC
GCCGT

#### 8.4 DNA sequences of PsCalA and PsCalB inserts in the pET28a(+) vector

The nucleotide sequences of the open reading frames (ORFs) for PsCalA and PsCalB inserted into the pET28a(+) vector are shown below. All other mutant enzymes or candidate constructs generated in this study were cloned using the identical strategy.

**PsCalA** insert (in pET28a(+)) nucleotide sequence:

ATCCGGATATAGTTCCTCCTTTCAGCAAAAAACCCCTCAAGACCCGTTTAGAGGCCCCAAGGGGTTATGCTAGTTAT
TGCTCAGCGGTGGCAGCAGCCAACTCAGCTTCCTTTCGGGCTTTGTTAGCAGCCGGATCTTCAAACGTAGGTAGACGCCA
GACCACCGTCAACCGGAATGTTGATACCGTTGATCCAACGAGATTCTTCCGCGCACATGAACGCGATAACCGCCGCAACT
TCGTCCGCATACGCCGGACGTTTGATACGATGCGCGTCCGCTTGGGTACGTTCCTGACCCAGCATAGTCACGAACTCGTT
CAGGATCGGGGTGAAAACTGGACCTGGCGCGATGCAGTTCATACGAACAGAGGTACGCATGAACCATTCCTGCGCCTGC
ACCTGAGACCAAACGATCAGCGCTTCTTTGAAGTACTGGTAGCAAAATTCCGGCGCAACCGGGTTCTGCTTCAGCCACGC
CTGACCTTCAGAGAAACCAACAACGGAACCCAGTTCTTTGTGCAGTTGCAGACGCGCTGGCCATTCAGCGCCCAGCACAG
AAGACACATTAACGATAGAACCACCCGGCTGGATACGAGAGAGAACCGCTTCGGTCAGGTATTTCAGACCCAGGTAGTTA
ACGTTCGCAACCAGCTGCGGGTCCGCGGTACCCGGAACACCCGCGATGTTGCACAGACCGTCGATCTTTTCCGGCAGCTG
AGAGATCGCTTTGTCGATACCCTCTGGGTGAGACAGGTCAGCCTGAACAAACGCGTCCAGGGTCAGAGACGGCATGTTA
CGGTCAACACCGATAACGGTCGCACCGTGAGAACGCAGAACACGCGCGGTTTCCGCACCAATACCAGAAGAAACACCGGT
AACAACAATCTTTTTGTTAGTGAGTTGCATGCCGCTGCTGTGATGATGATGATGATGGCTGCTGCCCATGGTATATCTCC
TTCTTAAAGTTAAACAAAATTATTTCTAGAGGGGAATTGTTATCCGCTCACAATTCCCCTATAGTGAGTCGTATTAATTT
CGCGGGATCGAGATCTCGATCCTCTACGCCGGACGCATCGTGGCCGGCATCACCGGCGCCACAGGTGCGGTTGCTGGCG
CCTATATCGCCGACATCACCGATGGGGAAGATCGGGCTCGCCACTTCGGGCTCATGAGCGCTTGTTTCGGCGTGGGTATG
GTGGCAGGCCCCGTGGCCGGGGGACTGTTGGGCGCCATCTCCTTGCATGCACCATTCCTTGCGGCGGCGGTGCTCAACG
GCCTCAACCTACTACTGGGCTGCTTCCTAATGCAGGAGTCGCATAAGGGAGAGCGTCGAGATCCCGGACACCATCGAATG
GCGCAAAACCTTTCGCGGTATGGCATGATAGCGCCCGGAAGAGAGTCAATTCAGGGTGGTGAATGTGAAACCAGTAACG
TTATACGATGTCGCAGAGTATGCCGGTGTCTCTTATCAGACCGTTTCCCGCGTGGTGAACCAGGCCAGCCACGTTTCTGC
GAAAACGCGGGAAAAAGTGGAAGCGGCGATGGCGGAGCTGAATTACATTCCCAACCGCGTGGCACAACAACTGGCGGGC
AAACAGTCGTTGCTGATTGGCGTTGCCACCTCCAGTCTGGCCCTGCACGCGCCGTCGCAAATTGTCGCGGCGATTAAATC
TCGCGCCGATCAACTGGGTGCCAGCGTGGTGGTGTCGATGGTAGAACGAAGCGGCGTCGAAGCCTGTAAAGCGGCGGTG
CACAATCTTCTCGCGCAACGCGTCAGTGGGCTGATCATTAACTATCCGCTGGATGACCAGGATGCCATTGCTGTGGAAGC
TGCCTGCACTAATGTTCCGGCGTTATTTCTTGATGTCTCTGACCAGACACCCATCAACAGTATTATTTTCTCCCATGAAG
ACGGTACGCGACTGGGCGTGGAGCATCTGGTCGCATTGGGTCACCAGCAAATCGCGCTGTTAGCGGGCCCATTAAGTTC
TGTCTCGGCGCGTCTGCGTCTGGCTGGCTGGCATAAATATCTCACTCGCAATCAAATTCAGCCGATAGCGGAACGGGAAG
GCGACTGGAGTGCCATGTCCGGTTTTCAACAAACCATGCAAATGCTGAATGAGGGCATCGTTCCCACTGCGATGCTGGTT
GCCAACGATCAGATGGCGCTGGGCGCAATGCGCGCCATTACCGAGTCCGGGCTGCGCGTTGGTGCGGATATCTCGGTAG
TGGGATACGACGATACCGAAGACAGCTCATGTTATATCCCGCCGTTAACCACCATCAAACAGGATTTTCGCCTGCTGGGG
CAAACCAGCGTGGACCGCTTGCTGCAACTCTCTCAGGGCCAGGCGGTGAAGGGCAATCAGCTGTTGCCCGTCTCACTGGT
GAAAAGAAAAACCACCCTGGCGCCCAATACGCAAACCGCCTCTCCCCGCGCGTTGGCCGATTCATTAATGCAGCTGGCAC
GACAGGTTTCCCGACTGGAAAGCGGGCAGTGAGCGCAACGCAATTAATGTAAGTTAGCTCACTCATTAGGCACCGGGATC
TCGACCGATGCCCTTGAGAGCCTTCAACCCAGTCAGCTCCTTCCGGTGGGCGCGGGGCATGACTATCGTCGCCGCACTTA
TGACTGTCTTCTTTATCATGCAACTCGTAGGACAGGTGCCGGCAGCGCTCTGGGTCATTTTCGGCGAGGACCGCTTTCGC
TGGAGCGCGACGATGATCGGCCTGTCGCTTGCGGTATTCGGAATCTTGCACGCCCTCGCTCAAGCCTTCGTCACTGGTCC
CGCCACCAAACGTTTCGGCGAGAAGCAGGCCATTATCGCCGGCATGGCGGCCCCACGGGTGCGCATGATCGTGCTCCTGT
CGTTGAGGACCCGGCTAGGCTGGCGGGGTTGCCTTACTGGTTAGCAGAATGAATCACCGATACGCGAGCGAACGTGAAG
CGACTGCTGCTGCAAAACGTCTGCGACCTGAGCAACAACATGAATGGTCTTCGGTTTCCGTGTTTCGTAAAGTCTGGAAA
CGCGGAAGTCAGCGCCCTGCACCATTATGTTCCGGATCTGCATCGCAGGATGCTGCTGGCTACCCTGTGGAACACCTACA

TCTGTATTAACGAAGCGCTGGCATTGACCCTGAGTGATTTTTCTCTGGTCCCGCCGCATCCATACCGCCAGTTGTTTACC
CTCACAACGTTCCAGTAACCGGGCATGTTCATCATCAGTAACCCGTATCGTGAGCATCCTCTCTCGTTTCATCGGTATCA
TTACCCCCATGAACAGAAATCCCCCTTACACGGAGGCATCAGTGACCAAACAGGAAAAAACCGCCCTTAACATGGCCCGC
TTTATCAGAAGCCAGACATTAACGCTTCTGGAGAAACTCAACGAGCTGGACGCGGATGAACAGGCAGACATCTGTGAATC
GCTTCACGACCACGCTGATGAGCTTTACCGCAGCTGCCTCGCGCGTTTCGGTGATGACGGTGAAAACCTCTGACACATGC
AGCTCCCGGAGACGGTCACAGCTTGTCTGTAAGCGGATGCCGGGAGCAGACAAGCCCGTCAGGGCGCGTCAGCGGGTGT
TGGCGGGTGTCGGGGCGCAGCCATGACCCAGTCACGTAGCGATAGCGGAGTGTATACTGGCTTAACTATGCGGCATCAG
AGCAGATTGTACTGAGAGTGCACCATATATGCGGTGTGAAATACCGCACAGATGCGTAAGGAGAAAATACCGCATCAGG
CGCTCTTCCGCTTCCTCGCTCACTGACTCGCTGCGCTCGGTCGTTCGGCTGCGGCGAGCGGTATCAGCTCACTCAAAGGC
GGTAATACGGTTATCCACAGAATCAGGGGATAACGCAGGAAAGAACATGTGAGCAAAAGGCCAGCAAAAGGCCAGGAAC
CGTAAAAAGGCCGCGTTGCTGGCGTTTTTCCATAGGCTCCGCCCCCCTGACGAGCATCACAAAAATCGACGCTCAAGTCA
GAGGTGGCGAAACCCGACAGGACTATAAAGATACCAGGCGTTTCCCCCTGGAAGCTCCCTCGTGCGCTCTCCTGTTCCGA
CCCTGCCGCTTACCGGATACCTGTCCGCCTTTCTCCCTTCGGGAAGCGTGGCGCTTTCTCATAGCTCACGCTGTAGGTAT
CTCAGTTCGGTGTAGGTCGTTCGCTCCAAGCTGGGCTGTGTGCACGAACCCCCCGTTCAGCCCGACCGCTGCGCCTTATC
CGGTAACTATCGTCTTGAGTCCAACCCGGTAAGACACGACTTATCGCCACTGGCAGCAGCCACTGGTAACAGGATTAGCA
GAGCGAGGTATGTAGGCGGTGCTACAGAGTTCTTGAAGTGGTGGCCTAACTACGGCTACACTAGAAGGACAGTATTTGG
TATCTGCGCTCTGCTGAAGCCAGTTACCTTCGGAAAAAGAGTTGGTAGCTCTTGATCCGGCAAACAAACCACCGCTGGTA
GCGGTGGTTTTTTTGTTTGCAAGCAGCAGATTACGCGCAGAAAAAAAGGATCTCAAGAAGATCCTTTGATCTTTTCTACG
GGGTCTGACGCTCAGTGGAACGAAAACTCACGTTAAGGGATTTTGGTCATGAACAATAAAACTGTCTGCTTACATAAACA
GTAATACAAGGGGTGTTATGAGCCATATTCAACGGGAAACGTCTTGCTCTAGGCCGCGATTAAATTCCAACATGGATGCT
GATTTATATGGGTATAAATGGGCTCGCGATAATGTCGGGCAATCAGGTGCGACAATCTATCGATTGTATGGGAAGCCCG
ATGCGCCAGAGTTGTTTCTGAAACATGGCAAAGGTAGCGTTGCCAATGATGTTACAGATGAGATGGTCAGACTAAACTG
GCTGACGGAATTTATGCCTCTTCCGACCATCAAGCATTTTATCCGTACTCCTGATGATGCATGGTTACTCACCACTGCGA
TCCCCGGGAAAACAGCATTCCAGGTATTAGAAGAATATCCTGATTCAGGTGAAAATATTGTTGATGCGCTGGCAGTGTTC
CTGCGCCGGTTGCATTCGATTCCTGTTTGTAATTGTCCTTTTAACAGCGATCGCGTATTTCGTCTCGCTCAGGCGCAATC
ACGAATGAATAACGGTTTGGTTGATGCGAGTGATTTTGATGACGAGCGTAATGGCTGGCCTGTTGAACAAGTCTGGAAA
GAAATGCATAAACTTTTGCCATTCTCACCGGATTCAGTCGTCACTCATGGTGATTTCTCACTTGATAACCTTATTTTTGAC
GAGGGGAAATTAATAGGTTGTATTGATGTTGGACGAGTCGGAATCGCAGACCGATACCAGGATCTTGCCATCCTATGGA
ACTGCCTCGGTGAGTTTTCTCCTTCATTACAGAAACGGCTTTTTCAAAAATATGGTATTGATAATCCTGATATGAATAAA
TTGCAGTTTCATTTGATGCTCGATGAGTTTTTCTAAGAATTAATTCATGAGCGGATACATATTTGAATGTATTTAGAAAA
ATAAACAAATAGGGGTTCCGCGCACATTTCCCCGAAAAGTGCCACCTGAAATTGTAAACGTTAATATTTTGTTAAAATTC
GCGTTAAATTTTTGTTAAATCAGCTCATTTTTTAACCAATAGGCCGAAATCGGCAAAATCCCTTATAAATCAAAAGAATA
GACCGAGATAGGGTTGAGTGTTGTTCCAGTTTGGAACAAGAGTCCACTATTAAAGAACGTGGACTCCAACGTCAAAGGG
CGAAAAACCGTCTATCAGGGCGATGGCCCACTACGTGAACCATCACCCTAATCAAGTTTTTTGGGGTCGAGGTGCCGTAA
AGCACTAAATCGGAACCCTAAAGGGAGCCCCCGATTTAGAGCTTGACGGGGAAAGCCGGCGAACGTGGCGAGAAAGGAA
GGGAAGAAAGCGAAAGGAGCGGGCGCTAGGGCGCTGGCAAGTGTAGCGGTCACGCTGCGCGTAACCACCACACCCGCCG
CGCTTAATGCGCCGCTACAGGGCGCGTCCCATTCGCCA

**PsCalB** insert (in pET28a(+)) nucleotide sequence:

ATCCGGATATAGTTCCTCCTTTCAGCAAAAAACCCCTCAAGACCCGTTTAGAGGCCCCAAGGGGTTATGCTAGTTAT
TGCTCAGCGGTGGCAGCAGCCAACTCAGCTTCCTTTCGGGCTTTGTTAGCAGCCGGATCTTCAACGGCGTGCCGCGCGGC
GGGCAAACGCCAGCATCGCGCCAATCAAGCGGCGGCGCGCGCCCCACGGCGGCCGGGCCAGACCGGTTGACGCCAGGCG
CGGCTGAACCATCACCGGGCGTTTTTTGCTAAAACGGGCAAAACCGTCAAAGCCGTGATATTCGCCGGTACCGGACGCGC
CAACGCCGCCAAACGGCAGATTCAGCTGCCCAACGTGGAACATGCAGTCGTTAAAGGTCACGCCGCCGCTCATGGTGCGG
GCAAGCACTTCATCCTGACGCGCGCCATCTTCGGCAAAGCAGTAAAGCGCCAGCGGGCGCGGGCGGGCATTCACAAACGC
CAGCGCATCATCCAGCGTTGGCGCAGGGATCACCGGCAGAATCGGGCCAAAAATTTCTTCGCGCATAAGCGCCGAATCTT
CCGGCGCATCAAACACCAGCATCGGCGGCATAAGGCGGCCGTTCGGCCCTGCAGGAACCGGCGCAAGTTCCACCACGCGC
GCGCCGCCTTCACGGGCTTCAGCCACCAAGCCTTTCAGGCGTTCCAAATGGCGGTCGCTGATAATGCGGGTGTAATCGCC
GTTCAAGCCGTTCGGGAACAACGCCGCAACCGCGCGGCGGGCTTCGGCTTCAAATTCAGCCATACGTGACGGCGGAACCA

ACGCATAATCCGGCGCAACGCAGGTCTGCCCGGCGTTCAGCAATTTGCCGGTCATCAGGCGCGGAACCGCGCGGCCCAGA
TCGGCACTATCATGAAGAATCGCCGGGGATTTACCGCCCAGTTCCAGGGTGACCGGCGTGAGGTTATCTGCCGCCGCGCG
CATCACTTCGCGGCCAACGCGGGTGGAGCCGGTAAAGACCAGATGATCAAGCGGCAGGCGGGTTAGCGCGCGCGCAAGC
GCCGCATCGCCGGTCACCACCGCCACCTGCCCTGGCGGGAATGCTTCTGCCACCAGCGCCGCCAGCGCTTCTGCGGTACG
CGGCGCCAGTTCGGACGGTTTAACCAGCACCCTGTTACCTGCCGCCAGCGCATCAACCATCGGCCCCAGGGTCAGCAGCA
CCGGATAGTTCCACGCGCCAATCACGCCGACCACGCCCAGCGGCTGATATTCATAAAACGCGCGCGCCGGCCAGAGAAAA
CGGCTACCCGCAACGCGGCGGCGGCGCATCCAGCCGCGAAGATGGCGGCGGGCATGGGCAATTTGGTCATAAAGCGGCA
GCAGTTCCAGCAGGCGGGTTTCTGCCGCCGTACGGCCGCCAAAATCGGCATCAACGGCGGCAATCAGGCCATCTTCATGG
CGGCCAAGCGCGCGGGCAAGCGCGGCAAGCGATTTGCGGCGGTCGGTCACATCCGGCCGCGGGCCTGCGGCATCAGACA
GATGGCGAAATGCCAGCGCAAGCGCGGGATGATCAGGATCAAGACTTGCGGTGGTTTGCGACATGCCGCTGCTGTGATG
ATGATGATGATGGCTGCTGCCCATGGTATATCTCCTTCTTAAAGTTAAACAAAATTATTTCTAGAGGGGAATTGTTATCC
GCTCACAATTCCCCTATAGTGAGTCGTATTAATTTCGCGGGATCGAGATCTCGATCCTCTACGCCGGACGCATCGTGGCC
GGCATCACCGGCGCCACAGGTGCGGTTGCTGGCGCCTATATCGCCGACATCACCGATGGGGAAGATCGGGCTCGCCACTT
CGGGCTCATGAGCGCTTGTTTCGGCGTGGGTATGGTGGCAGGCCCCGTGGCCGGGGGACTGTTGGGCGCCATCTCCTTG
CATGCACCATTCCTTGCGGCGGCGGTGCTCAACGGCCTCAACCTACTACTGGGCTGCTTCCTAATGCAGGAGTCGCATAA
GGGAGAGCGTCGAGATCCCGGACACCATCGAATGGCGCAAAACCTTTCGCGGTATGGCATGATAGCGCCCGGAAGAGAG
TCAATTCAGGGTGGTGAATGTGAAACCAGTAACGTTATACGATGTCGCAGAGTATGCCGGTGTCTCTTATCAGACCGTTT
CCCGCGTGGTGAACCAGGCCAGCCACGTTTCTGCGAAAACGCGGGAAAAAGTGGAAGCGGCGATGGCGGAGCTGAATTA
CATTCCCAACCGCGTGGCACAACAACTGGCGGGCAAACAGTCGTTGCTGATTGGCGTTGCCACCTCCAGTCTGGCCCTGC
ACGCGCCGTCGCAAATTGTCGCGGCGATTAAATCTCGCGCCGATCAACTGGGTGCCAGCGTGGTGGTGTCGATGGTAGA
ACGAAGCGGCGTCGAAGCCTGTAAAGCGGCGGTGCACAATCTTCTCGCGCAACGCGTCAGTGGGCTGATCATTAACTATC
CGCTGGATGACCAGGATGCCATTGCTGTGGAAGCTGCCTGCACTAATGTTCCGGCGTTATTTCTTGATGTCTCTGACCAG
ACACCCATCAACAGTATTATTTTCTCCCATGAAGACGGTACGCGACTGGGCGTGGAGCATCTGGTCGCATTGGGTCACCA
GCAAATCGCGCTGTTAGCGGGCCCATTAAGTTCTGTCTCGGCGCGTCTGCGTCTGGCTGGCTGGCATAAATATCTCACTC
GCAATCAAATTCAGCCGATAGCGGAACGGGAAGGCGACTGGAGTGCCATGTCCGGTTTTCAACAAACCATGCAAATGCTG
AATGAGGGCATCGTTCCCACTGCGATGCTGGTTGCCAACGATCAGATGGCGCTGGGCGCAATGCGCGCCATTACCGAGTC
CGGGCTGCGCGTTGGTGCGGATATCTCGGTAGTGGGATACGACGATACCGAAGACAGCTCATGTTATATCCCGCCGTTA
ACCACCATCAAACAGGATTTTCGCCTGCTGGGGCAAACCAGCGTGGACCGCTTGCTGCAACTCTCTCAGGGCCAGGCGGT
GAAGGGCAATCAGCTGTTGCCCGTCTCACTGGTGAAAAGAAAAACCACCCTGGCGCCCAATACGCAAACCGCCTCTCCCC
GCGCGTTGGCCGATTCATTAATGCAGCTGGCACGACAGGTTTCCCGACTGGAAAGCGGGCAGTGAGCGCAACGCAATTA
ATGTAAGTTAGCTCACTCATTAGGCACCGGGATCTCGACCGATGCCCTTGAGAGCCTTCAACCCAGTCAGCTCCTTCCGG
TGGGCGCGGGGCATGACTATCGTCGCCGCACTTATGACTGTCTTCTTTATCATGCAACTCGTAGGACAGGTGCCGGCAGC
GCTCTGGGTCATTTTCGGCGAGGACCGCTTTCGCTGGAGCGCGACGATGATCGGCCTGTCGCTTGCGGTATTCGGAATCT
TGCACGCCCTCGCTCAAGCCTTCGTCACTGGTCCCGCCACCAAACGTTTCGGCGAGAAGCAGGCCATTATCGCCGGCATG
GCGGCCCCACGGGTGCGCATGATCGTGCTCCTGTCGTTGAGGACCCGGCTAGGCTGGCGGGGTTGCCTTACTGGTTAGC
AGAATGAATCACCGATACGCGAGCGAACGTGAAGCGACTGCTGCTGCAAAACGTCTGCGACCTGAGCAACAACATGAATG
GTCTTCGGTTTCCGTGTTTCGTAAAGTCTGGAAACGCGGAAGTCAGCGCCCTGCACCATTATGTTCCGGATCTGCATCGC
AGGATGCTGCTGGCTACCCTGTGGAACACCTACATCTGTATTAACGAAGCGCTGGCATTGACCCTGAGTGATTTTTCTCT
GGTCCCGCCGCATCCATACCGCCAGTTGTTTACCCTCACAACGTTCCAGTAACCGGGCATGTTCATCATCAGTAACCCGT
ATCGTGAGCATCCTCTCTCGTTTCATCGGTATCATTACCCCCATGAACAGAAATCCCCCTTACACGGAGGCATCAGTGAC
CAAACAGGAAAAAACCGCCCTTAACATGGCCCGCTTTATCAGAAGCCAGACATTAACGCTTCTGGAGAAACTCAACGAGC
TGGACGCGGATGAACAGGCAGACATCTGTGAATCGCTTCACGACCACGCTGATGAGCTTTACCGCAGCTGCCTCGCGCGT
TTCGGTGATGACGGTGAAAACCTCTGACACATGCAGCTCCCGGAGACGGTCACAGCTTGTCTGTAAGCGGATGCCGGGA
GCAGACAAGCCCGTCAGGGCGCGTCAGCGGGTGTTGGCGGGTGTCGGGGCGCAGCCATGACCCAGTCACGTAGCGATAG
CGGAGTGTATACTGGCTTAACTATGCGGCATCAGAGCAGATTGTACTGAGAGTGCACCATATATGCGGTGTGAAATACCG
CACAGATGCGTAAGGAGAAAATACCGCATCAGGCGCTCTTCCGCTTCCTCGCTCACTGACTCGCTGCGCTCGGTCGTTCG
GCTGCGGCGAGCGGTATCAGCTCACTCAAAGGCGGTAATACGGTTATCCACAGAATCAGGGGATAACGCAGGAAAGAAC
ATGTGAGCAAAAGGCCAGCAAAAGGCCAGGAACCGTAAAAAGGCCGCGTTGCTGGCGTTTTTCCATAGGCTCCGCCCCCC

TGACGAGCATCACAAAAATCGACGCTCAAGTCAGAGGTGGCGAAACCCGACAGGACTATAAAGATACCAGGCGTTTCCCC
CTGGAAGCTCCCTCGTGCGCTCTCCTGTTCCGACCCTGCCGCTTACCGGATACCTGTCCGCCTTTCTCCCTTCGGGAAGC
GTGGCGCTTTCTCATAGCTCACGCTGTAGGTATCTCAGTTCGGTGTAGGTCGTTCGCTCCAAGCTGGGCTGTGTGCACGA
ACCCCCCGTTCAGCCCGACCGCTGCGCCTTATCCGGTAACTATCGTCTTGAGTCCAACCCGGTAAGACACGACTTATCGC
CACTGGCAGCAGCCACTGGTAACAGGATTAGCAGAGCGAGGTATGTAGGCGGTGCTACAGAGTTCTTGAAGTGGTGGCC
TAACTACGGCTACACTAGAAGGACAGTATTTGGTATCTGCGCTCTGCTGAAGCCAGTTACCTTCGGAAAAAGAGTTGGTA
GCTCTTGATCCGGCAAACAAACCACCGCTGGTAGCGGTGGTTTTTTTGTTTGCAAGCAGCAGATTACGCGCAGAAAAAAA
GGATCTCAAGAAGATCCTTTGATCTTTTCTACGGGGTCTGACGCTCAGTGGAACGAAAACTCACGTTAAGGGATTTTGGT
CATGAACAATAAAACTGTCTGCTTACATAAACAGTAATACAAGGGGTGTTATGAGCCATATTCAACGGGAAACGTCTTGC
TCTAGGCCGCGATTAAATTCCAACATGGATGCTGATTTATATGGGTATAAATGGGCTCGCGATAATGTCGGGCAATCAGG
TGCGACAATCTATCGATTGTATGGGAAGCCCGATGCGCCAGAGTTGTTTCTGAAACATGGCAAAGGTAGCGTTGCCAATG
ATGTTACAGATGAGATGGTCAGACTAAACTGGCTGACGGAATTTATGCCTCTTCCGACCATCAAGCATTTTATCCGTACT
CCTGATGATGCATGGTTACTCACCACTGCGATCCCCGGGAAAACAGCATTCCAGGTATTAGAAGAATATCCTGATTCAGG
TGAAAATATTGTTGATGCGCTGGCAGTGTTCCTGCGCCGGTTGCATTCGATTCCTGTTTGTAATTGTCCTTTTAACAGCG
ATCGCGTATTTCGTCTCGCTCAGGCGCAATCACGAATGAATAACGGTTTGGTTGATGCGAGTGATTTTGATGACGAGCGT
AATGGCTGGCCTGTTGAACAAGTCTGGAAAGAAATGCATAAACTTTTGCCATTCTCACCGGATTCAGTCGTCACTCATGG
TGATTTCTCACTTGATAACCTTATTTTTGACGAGGGGAAATTAATAGGTTGTATTGATGTTGGACGAGTCGGAATCGCAG
ACCGATACCAGGATCTTGCCATCCTATGGAACTGCCTCGGTGAGTTTTCTCCTTCATTACAGAAACGGCTTTTTCAAAAA
TATGGTATTGATAATCCTGATATGAATAAATTGCAGTTTCATTTGATGCTCGATGAGTTTTTCTAAGAATTAATTCATGA
GCGGATACATATTTGAATGTATTTAGAAAAATAAACAAATAGGGGTTCCGCGCACATTTCCCCGAAAAGTGCCACCTGAA
ATTGTAAACGTTAATATTTTGTTAAAATTCGCGTTAAATTTTTGTTAAATCAGCTCATTTTTTAACCAATAGGCCGAAAT
CGGCAAAATCCCTTATAAATCAAAAGAATAGACCGAGATAGGGTTGAGTGTTGTTCCAGTTTGGAACAAGAGTCCACTAT
TAAAGAACGTGGACTCCAACGTCAAAGGGCGAAAAACCGTCTATCAGGGCGATGGCCCACTACGTGAACCATCACCCTAA
TCAAGTTTTTTGGGGTCGAGGTGCCGTAAAGCACTAAATCGGAACCCTAAAGGGAGCCCCCGATTTAGAGCTTGACGGG
GAAAGCCGGCGAACGTGGCGAGAAAGGAAGGGAAGAAAGCGAAAGGAGCGGGCGCTAGGGCGCTGGCAAGTGTAGCGG
TCACGCTGCGCGTAACCACCACACCCGCCGCGCTTAATGCGCCGCTACAGGGCGCGTCCCATTCGCCA

### 9 Detailed results of wet-lab experiment validation

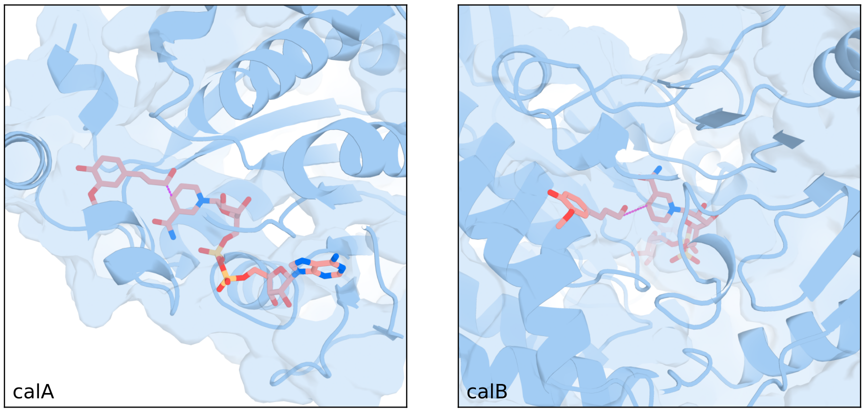


Figure S8: Schematic diagram of the binding pockets of CalA and CalB.

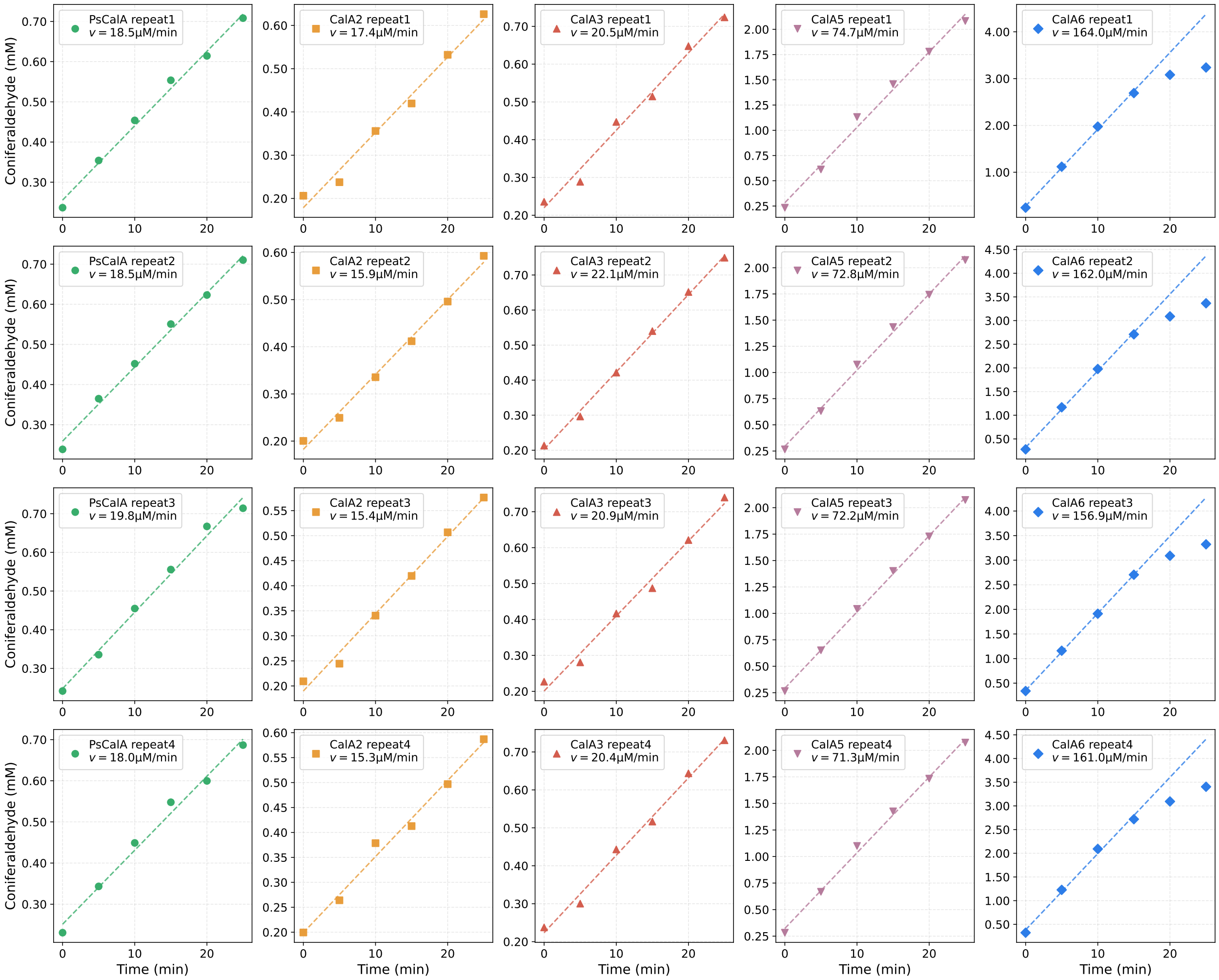


Figure S9: Reaction rates of purified enzymes PsCalA, CalA2, CalA3, CalA5, and CalA6.
